# The genetic consequences of range expansion and its influence on diploidization in polyploids

**DOI:** 10.1101/2023.10.18.562992

**Authors:** William W. Booker, Daniel R. Schrider

**Affiliations:** Department of Genetics, University of North Carolina at Chapel Hill, Chapel Hill, North Carolina, 27514-2916, United States of America

**Keywords:** Polyploid, population genetics, expansion load, allele surfing, diploidization, range expansion

## Abstract

Despite newly formed polyploids being subjected to myriad fitness consequences, the relative prevalence of polyploidy both contemporarily and in ancestral branches of the tree of life suggests alternative advantages that outweigh these consequences. One proposed advantage is that polyploids may more easily colonize novel habitats such as deglaciated areas. However, previous research conducted in diploids suggests that range expansion comes with a fitness cost as deleterious mutations may fix rapidly on the expansion front. Here, we interrogate the potential consequences of expansion in polyploids by conducting spatially explicit forward-in-time simulations to investigate how ploidy and inheritance patterns impact the relative ability of polyploids to expand their range. We show that under realistic dominance models, autopolyploids suffer greater fitness reductions than diploids as a result of range expansion due to the fixation of increased mutational load that is masked in the range core. Alternatively, the disomic inheritance of allopolyploids provides a shield to this fixation resulting in minimal fitness consequences. In light of this advantage provided by disomy, we investigate how range expansion may influence cytogenetic diploidization through the reversion to disomy in autotetraploids. We show that under a wide range of parameters investigated for two models of diploidization, disomy frequently evolves more rapidly on the expansion front than in the range core, and that this dynamic inheritance model has additional effects on fitness. Together our results point to a complex interaction between dominance, ploidy, inheritance, and recombination on fitness as a population spreads across a geographic range.

## INTRODUCTION

Polyploidy as a mechanism of evolution has had a profound influence on shaping the tree of life (Otto and Whitton 2000; Gregory and Mable 2005; Albertin and Marullo 2012; Barker et al. 2016; Zhan et al. 2016; Van de Peer et al. 2017; Li et al. 2018; One Thousand Plant Transcriptomes Initiative 2019). However, In many ways this observation does not reconcile with the known negative fitness consequences imbued by polyploidization. Once formed, polyploids often suffer meiotic abnormalities and genomic instability that result in reduced fertility and offspring viability as well as phenotypic instability (Mayer and Aguilera 1990; Comai et al. 2000; Ramsey and Schemske 2002; Morgan et al. 2020). Should a polyploid organism get past these hurdles, the minority cytotype exclusion principle suggests they have a very low probability of reproductive success by virtue of being surrounded by their more numerous non-polyploid relatives (Levin 1975; Husband 2000). Newly formed polyploids are also likely to be ecologically similar to their progenitor species, introducing additional pressures through the competition for resources (Rodriguez 1996; Yamauchi et al. 2004).

Despite these disadvantages, the prevalence of polyploid species suggests there are inherent advantages that polyploidy provides. One prevailing theory is that polyploidy provides an advantage in times of stress (Van de Peer et al. 2021), and they may more rapidly expand into novel or deglaciated habitats (Brochmann et al. 2004; Comai 2005; David 2022). The precise mechanisms that provide polyploids with these advantages aren’t entirely clear. One hypothesis proposes that polyploids simply have a greater adaptive potential resultant from a greater effective genic mutation rate (Otto 2007) and that duplicate genes allow for more specialized functionalization of one or both copies (Lynch and Conery 2000; Lynch and Force 2000; Adams and Wendel 2005; Gout and Lynch 2015). Similarly, more rapid adaptation may come from a multiplication of the gene regulatory network, as whole genome duplication creates a level of redundancy that frees parts of the network to reconfigure (Freeling and Thomas 2006; Hegarty and Hiscock 2007; Fusco et al. 2010; Yao et al. 2019), which can be particularly beneficial in times of environmental change (Ebadi et al. 2023). Alternatively, glacial cycles may play a more direct role in driving rates of polyploid formation by initiating secondary contact of diploid species (Stebbins 1984) potentially creating allopolyploids with fixed heterozygosity (Stebbins 1985; Brochmann et al. 2004).

The adaptive advantages provided by polyploidy for colonization of new habitats are not uncontroversial. Firstly, polyploids are not always more robust in harsher conditions and can have more restricted ranges (Levin 2002; Parisod et al. 2010). Although polyploids have a greater effective mutation rate that can produce novel adaptive mutations, this process also means polyploids will carry a greater mutational load that must be purged (Otto 2007). Additionally, the idea that mutations on duplicate genes are neutral because the non-mutated copy retains its function is not necessarily true (Mable and Otto 2001). Perhaps most glaring is the immediate reduction in effective population size due to the bottleneck resulting from polyploidization (Stebbins 1950), limiting the variation upon which selection can act for any adaptation.

Though there is a long history of discussion on the adaptive advantages polyploids may have in expanding their range, little work has been done to understand the genetic consequences of those expansions. As a species expands its range, populations along the expansion front are colonized by a small number of individuals that are more likely to colonize subsequent populations (Moreau et al. 2011). This phenomenon has a compounding effect, increasing an allele’s probability of fixation due to the elevated force of drift along the expansion front (Peischl et al. 2013; Peischl et al. 2015). This effect is particularly pronounced for recessive weakly deleterious mutations which can be masked from the effects of selection in larger core populations (Peischl and Excoffier 2015). As a result, populations on the edge of an expansion should have reduced fitness compared to populations in the core. Although the primary consequence of this process is reduced fitness, range expansions can alter the landscape of neutral genetic diversity across the genome (Schlichta et al. 2022) and can promote the hybridization of species coming into secondary contact (MacPherson et al. 2022). However, research on the consequences of range expansions have not been conducted in polyploids, and it is not currently known whether a geographically expanding polyploid population would experience stronger or milder consequences from a range expansion than an analogously expanding diploid population.

The ability for polyploids to mask deleterious alleles through whole genome duplication is often invoked to describe a relative advantage of polyploidy. However, theoretical work has shown that this masking results in a higher mutational load in polyploids (Haldane 1932; Hill 1970; Bever and Felber 1992) resulting in greater inbreeding depression in small populations and self-fertilizers (Bennett 1976). Because the mutational load present prior to expansion becomes a primary driver of expansion load (Peischl and Excoffier 2015), polyploids may be more severely impacted by the consequences of expanding their range. The effects of range expansion are also heavily influenced by recombination (Peischl et al. 2015)–a force that acts qualitatively differently during diploid and tetraploid meiosis (Grandont et al. 2013; Stenberg and Saura 2013), altering the ability of expanding migrants to purge the accumulating load. In total, these observations suggest that range expansion should have an inherently different effect on polyploid genomes.

Just as all polyploids must expand their range to ensure successful establishment, the ultimate fate of all polyploids is returning into a functionally diploid state (Wolfe 2001; Conant et al. 2014; Robertson et al. 2017; Mandáková and Lysak 2018; Li et al. 2021). While diploidization may refer to the process of fractionation (sometimes referred to as genic diploidization), where multiple copies of genes are generally lost or silenced (Freeling et al. 2012), diploidization first occurs cytogenetically to restore normal meiotic behavior. Because multivalent formation often results in meiotic errors, and therefore has large deleterious fitness effects, bivalent pairing is often restored relatively quickly (Feldman and Levy 2012; Feldman et al. 2012; Tayalé and Parisod 2013; Morgan et al. 2021). However, the fitness consequences of tetrasomic inheritance itself are less clear, and the restoration of disomy is often prolonged over millions of years while lineages continue to diversify (Lien et al. 2016; Scott et al. 2016; Robertson et al. 2017; Parey et al. 2022; Redmond et al. 2022). Because expansion load is driven by the fixation of deleterious recessive mutations, the evolution of disomy might then provide a fitness advantage to recently formed species by preventing the fixation of alleles across subgenomes.

Here, we explore both the genetic consequences of range expansion in polyploid species, and the role that these consequences play in driving cytogenetic diploidization and the evolution of disomy. Specifically, we develop a framework for spatially explicit forward-in-time simulations of autotetraploid, allotetraploid, and diploid species, simulating the expansion of species out of their core range. To evaluate the consequences of expansion alone, rather than additional consequences from adapting into novel or suboptimal environments, we conducted simulations in a constant selective environment. We investigate expansion load as it relates to the pattern of inheritance, ploidy, and dominance to better understand how the range expansion of polyploids contributes to their fitness in comparison to diploids. In addition, we consider how selection for disomic inheritance may be influenced by expansion load by developing two models of diploidization and investigating the parameters in which diploidization is favored or disfavored in a geographically expanding species. Ultimately, we aim to identify the scenarios in which polyploidy and diploidization provide advantages or disadvantages for expansion and therefore contribute to their probability of establishment in novel regions.

## METHODS

### Simulation of Tetraploids

All simulations were conducted in SLiM version 4.0.1 (Haller and Messer 2023). Although SLiM does not support polyploid species natively, modeling polyploids is possible through some modifications. In general, these modifications were to the behavior of recombination and segregation and to fitness calculations in polyploid individuals.

To modify recombination for tetraploids, we added an additional population to SLiM that was reproductively isolated from all other populations and that served as storage for the additional two chromosomes present in each tetraploid individual. During initialization, each individual in the storage population is assigned, or “tagged”, to an individual in the focal population, resulting in the unique association of four chromosomes with each individual in the focal population such that these chromosomes could be accessed for reproduction and fitness calculations. For all substantive aspects of the simulation, the storage population served only to store the additional chromosomes used for modeling the individuals in the focal populations. For both auto- and allopolyploids we assume all chromosomes form bivalents and there is no double reduction. During reproduction, for autopolyploids we model tetrasomic inheritance by, for each parent, choosing one of three possible pairing combinations at random and forming a single recombinant chromosome within each pair resulting in two chromosomes passed on to a new offspring for that parent. This process is repeated with the second parent, and all chromosomes are tagged and separated into the storage and focal individuals (Supp. Fig. 1a). For allopolyploids, disomic inheritance was modeled by limiting pairing to the chromosomes within the storage and focal populations separately: two recombinant chromosomes from the focal population (one from each parent) were given to an individual in the focal population corresponding to the new offspring; similarly, recombinant chromosomes from the storage population for each parent were given to the an individual in the storage population corresponding to the second subgenome for this offspring individual (Supp. Fig. 1b).

To calculate fitness for tetraploids, first the count of derived alleles across all four chromosomes was obtained. This allele count was then used to obtain the appropriate dominance coefficient from the *h* vector (Table 1) or Eq. 3 below, and this dominance coefficient was then used in the fitness calculation in Eq. 2. Although diploids only have a single dominance coefficient given a single heterozygous state, tetraploids have three coefficients, corresponding to the possible heterozygote copy numbers of the derived allele (one, two, or three). As such, the vector of dominance coefficients depends on the model of dominance under consideration. All dominance coefficient vectors, excluding those obtained from the *h-s* relationship model (described below), can be found in Table 1. To determine the effect of expansion by itself, all fitnesses were rescaled to one by dividing each population’s fitness by the fitness of the outermost burn-in population at the start of expansion.

**Table 1.** Dominance coefficients (*h*) for deleterious mutations in diploids and tetraploids for each model and allelic state.

| Model | Diploid $h$ vector ($h_0, h_1, h_2$) | Tetraploid $h$ vector ($h_0, h_1, h_2, h_3, h_4$) |
| --- | --- | --- |
| Additive | (0.0, 0.5, 1.0) | (0.0, 0.25, 0.50, 0.75, 1.0) |
| Recessive | (0.0, 0.0, 1.0) | (0.0, 0.0, 0.0, 0.0, 1.0) |
| Recessive (duplex) | (0.0, 0.0, 1.0) | (0.0, 0.0, 0.0, 1.0, 1.0) |

### Demographic Model

We used a stepping stone model where the total population consisted of a series of demes along a single dimension, and migration occurred between adjacent demes. Prior to expansion, demes at both edges were reflecting, with no migrants expanding beyond either edge. During expansion, reflection at one edge was removed, and migration beyond this deme resulted in the colonization of an additional deme.

Within each deme, we allowed population size to grow according to the logistic growth model adapted from Peischl et al. (2013) and defined in Eq. 1:

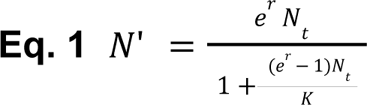

Here, *N_t_* is the number of individuals at time *t,* and *N’* is the rate parameter of a poisson distribution representing the number of individuals at time *t* + 1. *K* is the carrying capacity of the deme and *r* is the intrinsic growth rate. For all simulations, generations are discrete and non-overlapping. At the start of each generation, individuals are killed probabilistically as a function of their fitness (P = 1-*w*), and reproduction immediately follows this step. Individuals are monecious, and mates are chosen at random with replacement until *N_t_*_+1_ ∼ Poisson(*N’*) offspring are generated. We draw the number of offspring from a Poisson distribution to add stochastic dynamics present in natural populations, which can also lead to the extinction of individual demes. Individual fitness has no effect on reproductive success among survivors of the death phase. Following reproduction, individuals migrated to adjacent populations at rate *m*, with the direction of migration chosen at random, provided individuals were not in the first or final demes during burn-in.

### Model Parameters

At the beginning of each simulation replicate, 5 demes were initialized at carrying capacity and allowed to evolve for a burn-in of 5000 generations, and migration was limited to between adjacent demes among these 5 initial demes only. We chose 10*N* generations (5 demes at *N*=100; 5000 generations) as the burn-in because across all models, nucleotide diversity plateaus prior to or approximately at 10*N* generations (see Supp. Fig. 2). Following this burn-in, individuals were allowed to migrate into new demes from the edge deme, and at this point migration was allowed between all pairs of adjacent demes. For all simulations, the carrying capacity *K* was set to 100, *r*=log(2), *m*=0.05. For the genomic parameters, each individual had a 1 Mb genome, deleterious mutations occurred at a rate of 2.5×10^-8^, and beneficial mutations occurred at a rate of 5×10^-9^. For all simulations the recombination rate (⍴) was set to 1.0×10^-6^ resulting in approximately 1 recombination event per chromosome per generation.

In this model, selection is density independent as mutations do not affect reproductive output, and the impact of mutations on survival is not dependent on the population density (see Travis et al. 2013). Fitness only affects survival, and was calculated multiplicatively as shown in Eq. 2, where *w* is the individual’s fitness and *s* is the selection coefficient for the given mutation *i*, and *h_j_* is the dominance coefficient for mutation *i* given that it is present at copy number *j* in the individual under consideration:

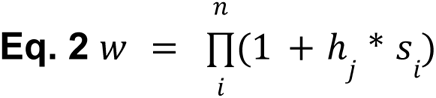

For all simulations, the selection coefficient *s* for deleterious mutations was drawn from a gamma distribution equal parameterized with *α* = 0.16 and *β* = 0.0092 (mean = -0.001472), taken from Huber et al. (2018). All beneficial mutations had a fixed *s* of the deleterious gamma mean, 0.001472. To calculate the effect of mutations outside of the homozygous state, we multiplied *s* by the dominance coefficient for the individual’s copy number of the derived allele as specified by the dominance model being simulated (Table 1). Here, the additive model describes a DFE where the fitness effect scales linearly with the proportion of alleles in the mutated state. The recessive model describes a DFE where there is no selective effect for deleterious mutations as long as one wild-type allele is present at the locus, and the duplex model describes a DFE where selection is dosage dependent and acts only when the wild-type allele is lower than 50% in frequency for deleterious mutations (see Hill 1970). For the additive model, the *h* vector for beneficial mutations was equal to that for deleterious mutations. For all other models beneficial mutations were treated as dominant.

### Modeling an empirical DFE specifying a relationship between the selection and dominance coefficients

In an attempt to model the effects of expansion load on more realistic populations, we modeled an expanding population experiencing mutations whose fitness effects were drawn from an empirically estimated DFE. To do so, we used the genome wide *Arabidopsis lyrata* DFE from Huber et al. (2018). In addition to having a distribution of selection coefficients, the model from Huber et al. (2018) also modeled the relationship between dominance coefficient *h* and *s* (Eq. 3). Here, *θ*_intercept_ defines the value of *h* at s = 0, and *θ*_rate_ defines the rate at which *h* approaches 0 with a declining negative *s*. To broadly summarize the distributions of *s* and *h* defined by this model, all mutations are deleterious, with most being weakly deleterious, and there is a negative relationship between *h* and *s* such that strongly deleterious mutations are more recessive. For our simulations, the DFE was parameterized with *s* as a gamma distribution with *α* = 0.16 and *β* = 0.0092, multiplied by -1 to make mutations deleterious, and *h* was parameterized with *θ*_intercept_ = 0.978 *θ*_rate_ = 50328. Importantly, the *s* in Eq. 3 is negative, ensuring all *h* values are between 0 and 1.

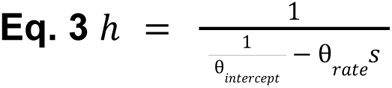

To translate the dominance coefficient from the diploid *h* to tetraploids, we used the flux model of dominance from Kascer and Burns (1981) by setting a mutation’s *h* value obtained from Eq. 3 as the flux value at 0.5 wild-type frequency. Briefly, this model describes a non-linear relationship between enzymatic activity regulated by the number of wild-type allelic copies, and the output of a system (flux) which is the phenotypic effect. Under this model, the phenotype value increases continuously with the input (originally the amount of enzyme activity), allowing the translation from a diploid to tetraploid model by using an individual’s fraction of non-mutant alleles at a site as the input (0, 0.5, or 1 for diploids, and 0, 0.25, 0.5, 0.75, or 1 for tetraploids), denoted by *P* in the equations below. Eq. 4 is adapted from Boucher et al. (2016) and models the shape of the flux curve. Here, *W* is the functional output of a system, which we treat as fitness rather than flux. *W* is determined by the proportion of ancestral or wild-type alleles *P*, according to a curve whose shape is determined by the coefficient *B*, which is in turn determined by the diploid dominance coefficient *h*. Because the output of the system at *P* = 0.5 is equal to 1 - *h*, we can use Eq. 5 to solve for *B* given any *h*, shown in Eq. 6. Combining equations 4 and 6, we can then solve for the output of the system given *P* and *h*, shown in Eq. 7. Because we are interested in the fitness consequences of a mutation given its copy number in an individual, and *h* only describes the fitness effects at 50% frequency, we then calculate *Z*=1-*W*, giving us the total fitness cost for a mutation given its frequency *P* and *h*, shown in Eq. 8. Importantly, Eqs. 6-8 are not solvable for *h* = 0.5, however the curve at this value represents the additive model therefore the desired dominance coefficient *Z* in such cases can be trivially obtained from *P* (see Supp. Fig. 3).

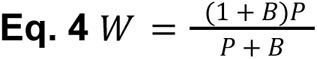

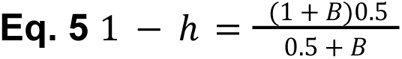

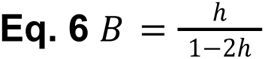

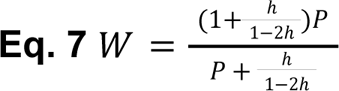

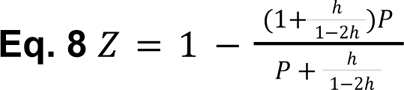

### Cytogenetic Diploidization

To model the evolution of cytogenetic diploidization in range-expanding populations, we ran our simulations as described previously, but allowed for additional mutations that altered the inheritance pattern through the preferential pairing of chromosomes. All individuals start as autopolyploids with bivalent pairing but random assortment between all four chromosomes. As diploidization mutations are added, the pairing behavior of chromosomes is altered depending on the model of diploidization being simulated (see below). Importantly, for these mutations we did not assume an infinite sites model, and allowed diploidization mutations to revert to the ancestral state.

Although the mechanisms for diploidization are not entirely clear, evidence suggests two distinct processes are largely responsible (see Li et al. 2021). One process involves mutations that regulate the meiotic machinery which fully or partially restore normal meiotic pairing behavior (Sánchez-Morán et al. 2001; Jenczewski et al. 2003; Liu et al. 2006; Henry et al. 2014; Gonzalo et al. 2019; Morgan et al. 2020). We refer to this as the meiotic gene model, and we define its parameters in Eq. 9. Here, *d* is the number of unique derived diploidization alleles in an individual, and *λ* is the number of mutations required to restore disomic inheritance. Under this model, subgenomes are predefined but all chromosomes freely recombine. As diploidization mutations occur, each mutation has a fixed effect in weighting the probability of recombination between chromosomes of the same subgenome, until 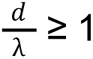, at which point the subgenomes are differentiated and never recombine. Although the process of diploidization via this model is not well understood, our purpose for modeling cytogenetic diploidization in this fashion is to determine whether the presence of some mutation(s) of large effect on disomic behavior, which may provide an adaptive benefit to an expanding population, was more or less likely to result in diploidization as a result of that expansion. For all models investigated, we varied the recombination rate (⍴), selective coefficient (*s*), and the number of mutations required to restore disomy (*λ*). The rate at which diploidization mutations arose (*μ_dip_*) was set to 2.5×10^-10^ multiplied by *λ* (e.g. *μ_dip_* = 5×10^-10^ for *λ* = 2) such that diploidization occurred at a similar rate across all models.

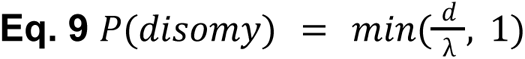

Alternatively, diploidization may occur as a process of chromosomal differentiation, where large rearrangements, duplications, deletions, or other structural changes drive differences between chromosomes such that they are no longer able to pair. Le Comber et al. (2010) showed that drift of neutral mutations resulting in chromosomal differences alongside fitness consequences for pairing failures was sufficient to drive the evolution of diploidization. Because expansion is akin to accelerating the force of drift, it is possible expansion may more rapidly result in the evolution of diploidization.

To model this behavior, we adapted the model from Le Comber et al. (2010) and applied it to our simulations. Le Comber et al. (2010) describe a model in which, as chromosomes diverge, their probability of pairing becomes less likely, and there are fitness costs associated with poor pairing behavior. This behavior is described by a sigmoid function (shown in Supp. Fig. 4), whereby pairing efficiency begins to decline at around 90% similarity between chromosomes, with similarity below 80% resulting in very poor pairing efficiency (approximately 0) and above 90% having essentially perfect efficiency (approximately 1). The steep slope of the curve for this model was based on observations in autotetraploid rye (Jenkins and Chatterjee 1994), and the 80% threshold where pairing drops to ∼0 was chosen as a compromise between experimental studies showing high variability in the conditions where recombination is suppressed (Okumura et al. 1987; Opperman et al. 2004). Because this threshold and slope likely have a significant influence on the behavior of this model, we described pairing efficiency using a more general sigmoid function to be able to alter these parameters, shown in Eq. 10.

Under this model, we allowed for *λ* diploidization loci randomly distributed across the chromosome to evolve at a rate of *μ_dip_* mutations per locus per generation. Diploidization mutations have no fitness effects themselves and only affect the pairing efficiency (*P*(pairing*_ij_*)). At the start of reproduction, the percent sequence identity is calculated using the difference in diploidization mutations between chromosomes *i* and *j*, *d_ij_*,for each combination of chromosome pairs. Pairing efficiency is then calculated with δ determining the slope and *β* determining the inflection point of the sigmoid curve. Once pairing efficiency is calculated for each combination of chromosomes, the average efficiency for each possible set of pairs (i.e. ab-cd, ac-bd, ad-bc) is calculated, and the set of pairs is chosen by random sample weighted by their pairing efficiency. This process is completed for each parent, and the probability of successful reproduction is calculated based on the average pairing efficiency of the two parents. Should the offspring not survive, another random set of parents is chosen to reproduce, and the process is repeated until a successful offspring is produced. The intention here is to mimic errors in meiosis resulting in inviable offspring as a result of improper chromosome pairing (referred below as pairing associated fitness costs, or PAFCs), a phenomenon common to neo-polyploids (Mayer and Aguilera 1990; Grandont et al. 2013; Bomblies et al. 2016; Morgan et al. 2020). Specifically, this models behavior such that if chromosomes are too differentiated to properly pair during meiosis, due to either one or more chromosomes being too differentiated from all others (under tetrasomy) or there being too much differentiation within subgenomes (under disomy), the resulting gametes from meiosis will be more likely to have an irregular number of chromosomes (i.e. aneuploidy) and thus may not be viable.

We investigated the evolution of disomy under this model using three combinations of values for parameters δ and *β* (Supp. Fig. 4). To mimic the behavior in Le Comber et al. (2011), with a steep slope and threshold ∼20% divergence, we used δ = 1 and *β* = 85. To determine the effect of this threshold, we also set the δ = 1 and *β* = 70 such that the decrease from 1 to 0 efficiency occurs at the same rate, but the pairing efficiency decline occurs at a greater sequence divergence. As well, we altered both parameters to δ = 0.2 and *β* = 85 to model a more gradual slope where the inflection point of pairing efficiency is at the same level of divergence as in the first model, but 0 efficiency is achieved at a much greater degree of divergence than in the Le Comber et al. (2011) model. Additionally, we tested these parameter combinations under two values of *λ*, 100 and 1000, and *μ_dip_* was set to 2×10^-3^ for *λ* = 100 and 1×10^-3^ for *λ* = 1000. Final values of *μ_dip_* were determined through extensive testing to ensure diploidization occurred in some demes within 1000 generations under a wide variety of parameter combinations. Similar to Le Comber et al. (2010) the mutation rate (i.e. *μ_dip_* ) did not affect the nature of diploidization spatially and only affected the rate at which diploidization was achieved, unless this rate was arbitrarily high causing rapid fluctuations in pairing efficiency.

To determine if disomy had evolved, we calculated the diploidization index as the difference in pairing efficiency between the top two possible sets of pairs, averaged across all individuals undergoing gametogenesis in a given generation. The rationale behind the diploidization index is that under disomy one possible pair should be at near 1 pairing efficiency and all other pairs at 0.

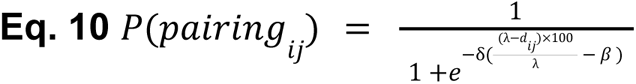

## GLOSSARY PRIMER

### The terms in this glossary are defined for the purposes of this paper and are not meant as general biological definitions

<u>Allopolyploid</u> - Organism with a duplicated genome defined here as having disomic inheritance and distinct subgenomes where no recombination occurs between them

<u>Autopolyploid</u> - Organism with a duplicated genome defined here as having tetrasomic inheritance and non-distinct subgenomes where recombination can occur between any pair of chromosome copies

<u>Neo-polyploid</u> - Polyploid that has recently originated and where no loss, silencing, or subfunctionalization of genes has occurred, and in the case of autopolyploids, no cytogenetic diploidization has occurred

<u>Cytogenetic diploidization</u> - The process of a polyploid transitioning from tetrasomic inheritance to disomic inheritance

<u>Disomic inheritance</u> - Chromosomal inheritance where preferential pairing between chromosomes occurs and the frequencies of allelic combinations in gametes are unequal, with two of the six possible combinations from alleles *A_1_A_2_A_3_A_4_* from a tetraploid parent absent

<u>Tetrasomic inheritance</u> - Chromosomal inheritance where no preferential pairing between chromosomes occurs and all allelic combinations in gametes are produced in equal frequencies

<u>Multivalent</u> - 3 or more chromosomes that associate and form multiple chiasmata across more than one chromosome pair during meiosis

<u>Bivalent</u> - A single pair of chromosomes associating and recombining during meiosis

<u>Expansion Load</u> - The buildup of deleterious mutations on the edge of the spatial distribution of an expanding species as a result of that expansion

## RESULTS

### Expansion load has a similar effect across tetraploids and diploids for additive models

To examine the consequences of expansion load in polyploids without the effects of dominance, we first modeled populations expanding their range with mutations having an additive effect on fitness relative to their frequency. Under this additive model, the reduction in fitness in the edge of the species range as a result of geographic expansion is remarkably similar across all tetraploid and diploid models, although autotetraploids may have slightly higher fitness along the edge (Fig. 1a; Fig. 2a). Interestingly, allotetraploids have a much worse starting unscaled fitness but experience a linear decline in edge fitness as the expansion proceeds that has a very similar slope to that of autopolyploids and diploids (Supp. Fig. 5). Core fitness declines in allotetraploids while increasing in autotetraploids and diploids at comparable rates, (Supp. Fig. 6a). Allopolyploids have a greater effective mutation rate than diploids, but do not benefit from a greater effective recombination rate–making deleterious mutations appear more rapidly but without any concomitant improvement in the ability to purge them.

**Figure 1.**
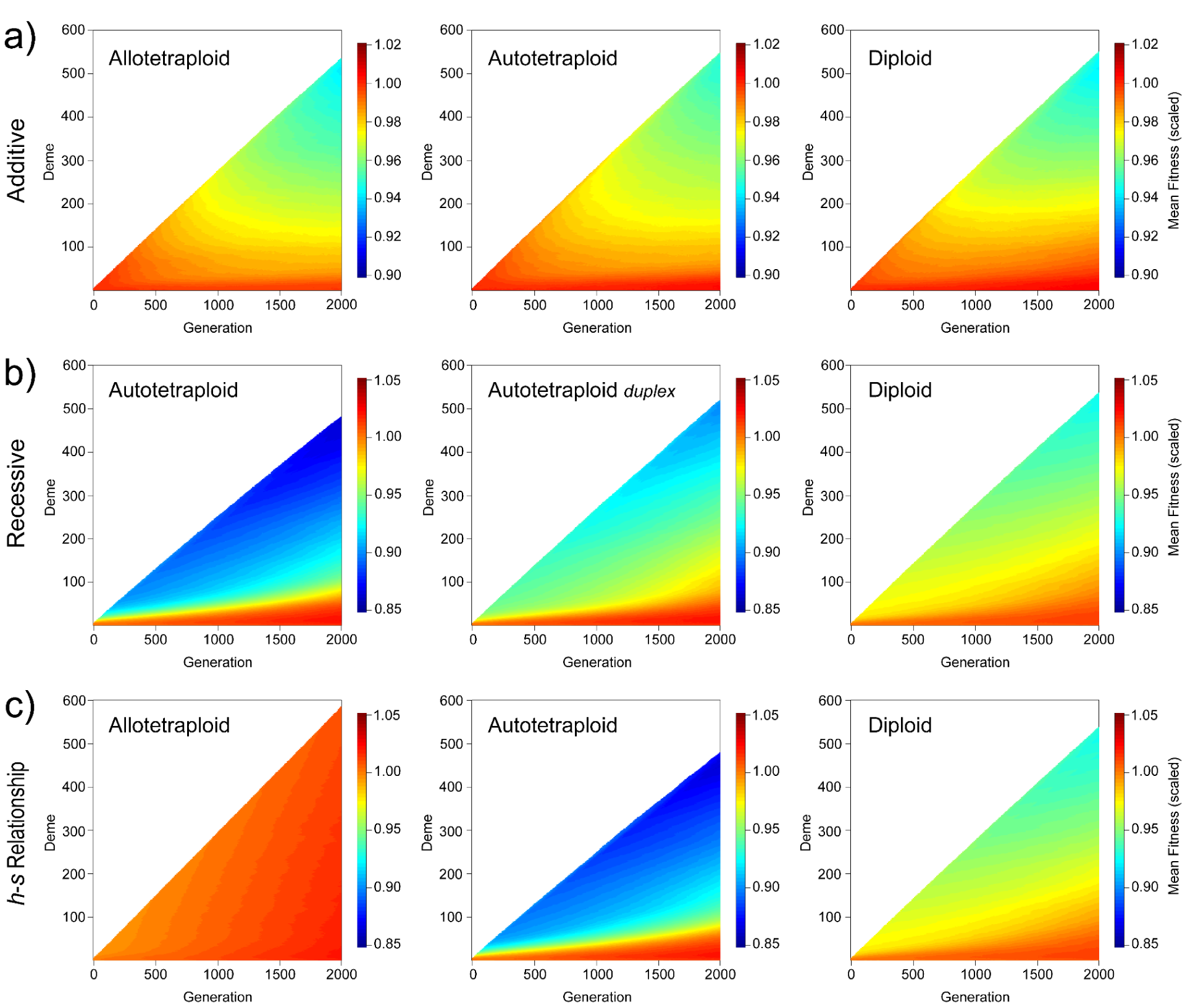
Mean fitness in each deme at each generation under additive (a), recessive (b), and *h*-*s* relationship (c) dominance models. Demes are only shown if 75% of replicates had that deme occupied to reduce artifacts from higher-fitness replicates expanding more quickly. Fitnesses for each deme are scaled by the initial fitness of the core at the onset of the range expansion, and then averaged across all replicates. Note that the scales are different for each dominance model.

**Figure 2.**
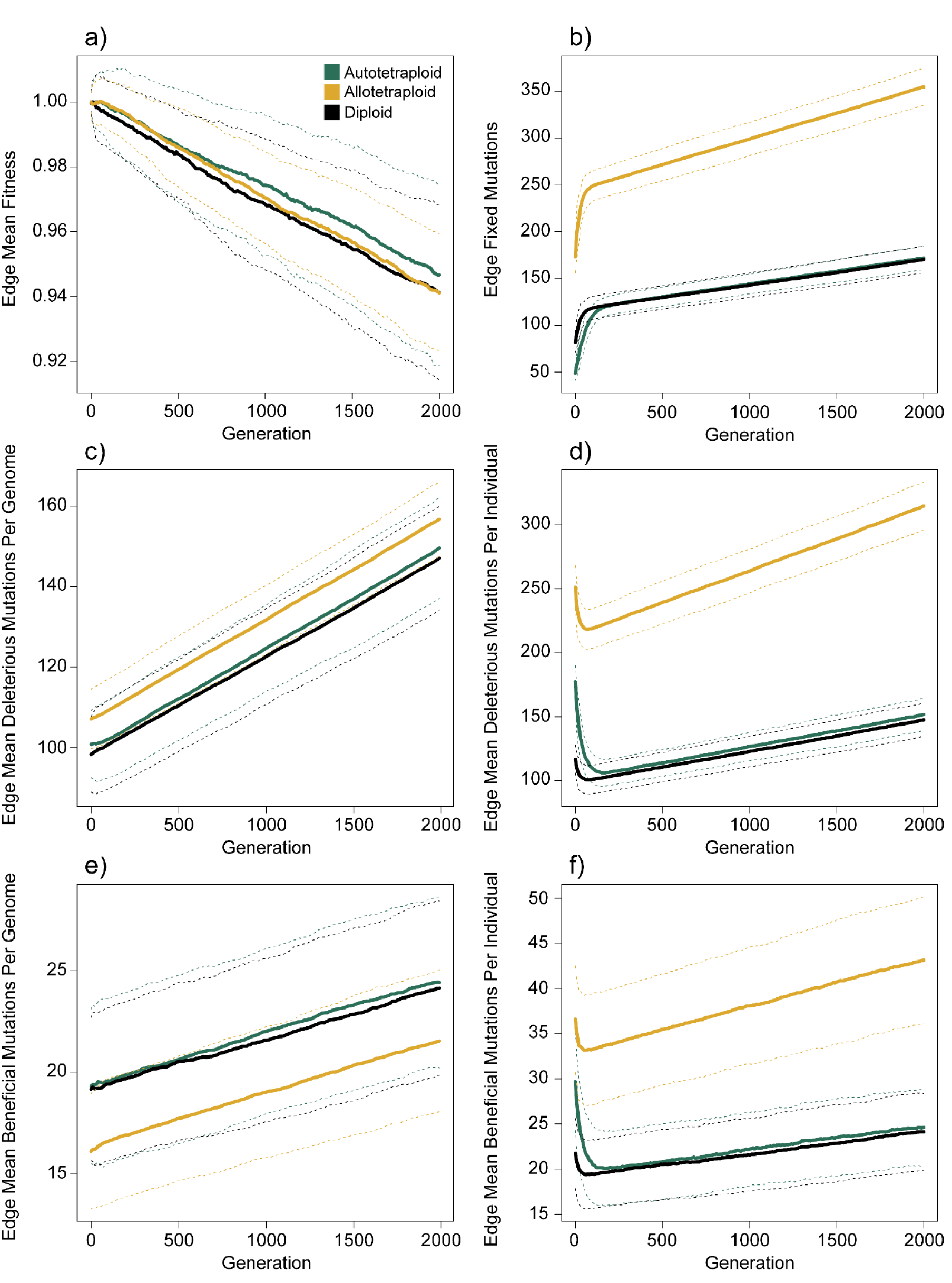
Statistics summarizing fitness and variation within spatially expanding populations under the additive dominance model. (a) Mean fitness (scaled) of the edge deme. (b) Number of fixed mutations in the edge deme. For allopolyploids, fixed mutations are those fixed within each subgenome. (c) Mean number of deleterious mutations per genome in the outermost deme (i.e. the edge). (d) Mean number of deleterious mutations per individual in the edge deme. (e) Mean number of beneficial mutations per genome in the outermost deme (i.e. the edge). (f) Mean number of beneficial mutations per individual in the edge deme. Lines show the average value of each statistic across 25 replicates, and dashed lines show ± 1 standard deviation.

Autopolyploids also have a greater effective mutation rate, but also allow for recombination between all possible pairs of four chromosomes which reduces the probability that mutations reach high frequency. As such, autotetraploids start with a generally higher number of deleterious mutations per individual than diploids (Fig. 2d), but a similar number of deleterious mutations per genome (Fig. 2c), suggesting that many of these mutations are at low frequency. This in turn results in a large fraction of mutations in tetraploids to be lost quickly at the start of expansion before later increasing linearly over time at a similar rate as for diploids (Fig. 2d), with most of this increase attributed to fixed mutations (compare Fig. 2b,d).

### The masking effect of tetraploidy in core populations results in rapid fitness decline during expansion under recessive models

We next examined the consequences of expansion load in polyploids under a more realistic model where all deleterious mutations are recessive. Under both the fully recessive and duplex models, where selection acts if mutations are homozygous or have fewer than two copies of the wild-type allele, respectively, fitness in autotetraploids decreases rapidly on the expansion front (Fig. 1b). The magnitude of this fitness decline in the early stages of expansion in autotetraploids is significantly greater than that observed in diploids, although the decline eventually steadies out to a similar slope in both ploidies (Fig. 3a). A further look at the dynamics driving this pattern demonstrate that autopolyploid genomes and individuals have more deleterious mutations at the start of the expansion (Fig. 3c,d) resulting in a much greater number of deleterious mutations that fix along the expansion front (Fig. 3b). Finally, In contrast to the edge, fitness in the core increases at a faster rate in autopolyploid populations for both fully recessive and duplex models (Supp. Fig. 6b).

**Figure 3.**
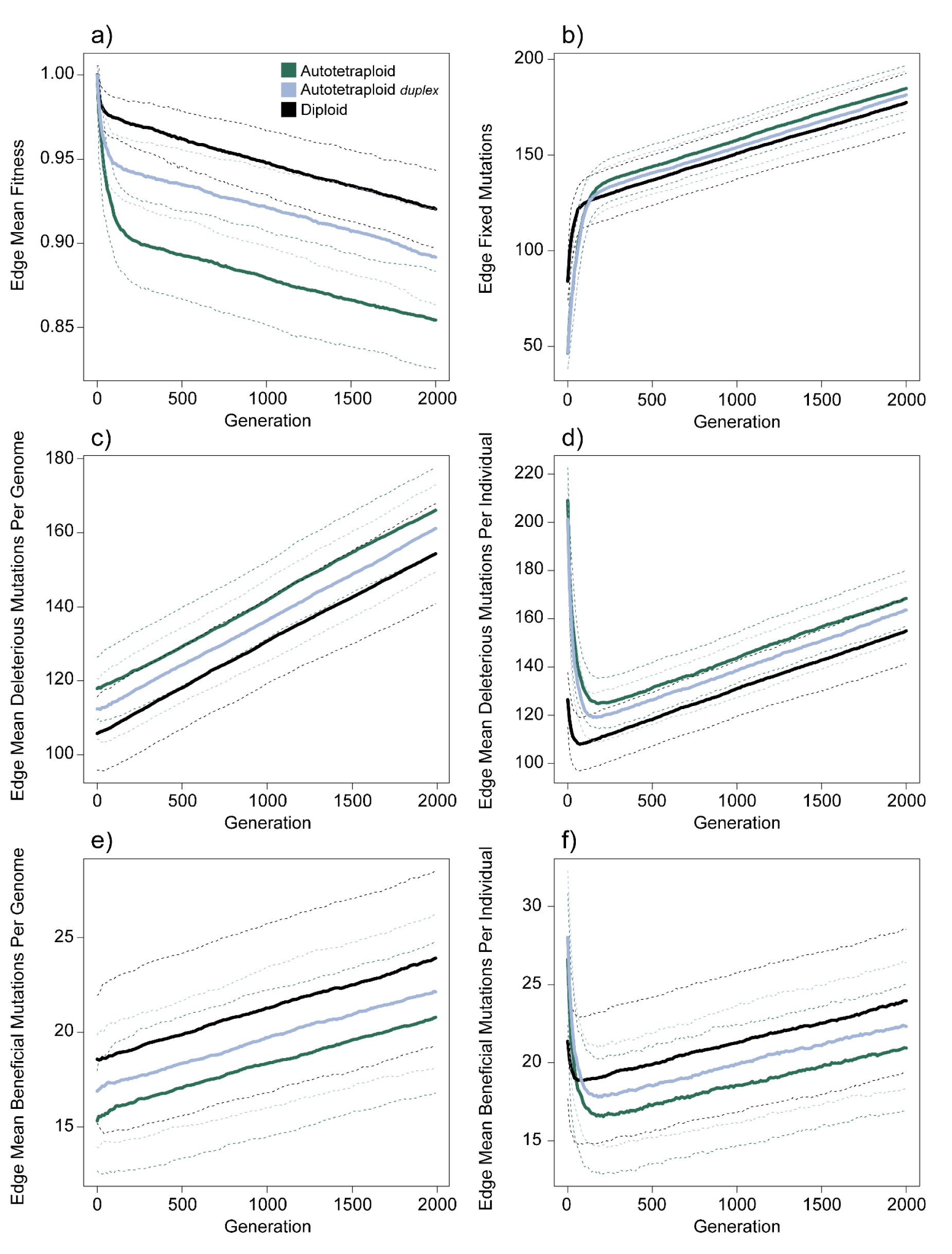
Statistics summarizing fitness and variation within spatially expanding populations under the recessive model. (a) Mean fitness (scaled) of the edge deme. (b) Number of fixed mutations in the edge deme. For allopolyploids, fixed mutations are those fixed within each subgenome. (c) Mean number of deleterious mutations per genome in the outermost deme (i.e. the edge). (d) Mean number of deleterious mutations per individual in the edge deme. (e) Mean number of beneficial mutations per genome in the outermost deme (i.e. the edge). (f) Mean number of beneficial mutations per individual in the edge deme. Lines show the average value of each statistic across 100 replicates, and dashed lines show ± 1 standard deviation.

### Autotetraploids behave similarly under an empirically estimated h-s relationship model as in the recessive model

We next examined a model that was adapted from one estimated in *Arabidopsis lyrata* (Huber et al. 2018) that models the relationship between the dominance and selection coefficients of deleterious mutations (Methods). Under this *h-s* relationship model, autotetraploids and diploids behave generally similarly to the fully recessive model (Fig. 1c, Fig. 4a-f). Again, autotetraploids accumulate more deleterious mutations during the burn-in because most mutations in the DFE are weakly deleterious and therefore proliferate in an autotetraploid where they are generally masked from the effects of selection, and as in the recessive model, exhibit an initial rapid drop prior to linear growth in the number of deleterious mutations. Also similar to the recessive model, in the core populations autotetraploid fitness exceeds diploid fitness (Supp Fig. 6c).

**Figure 4.**
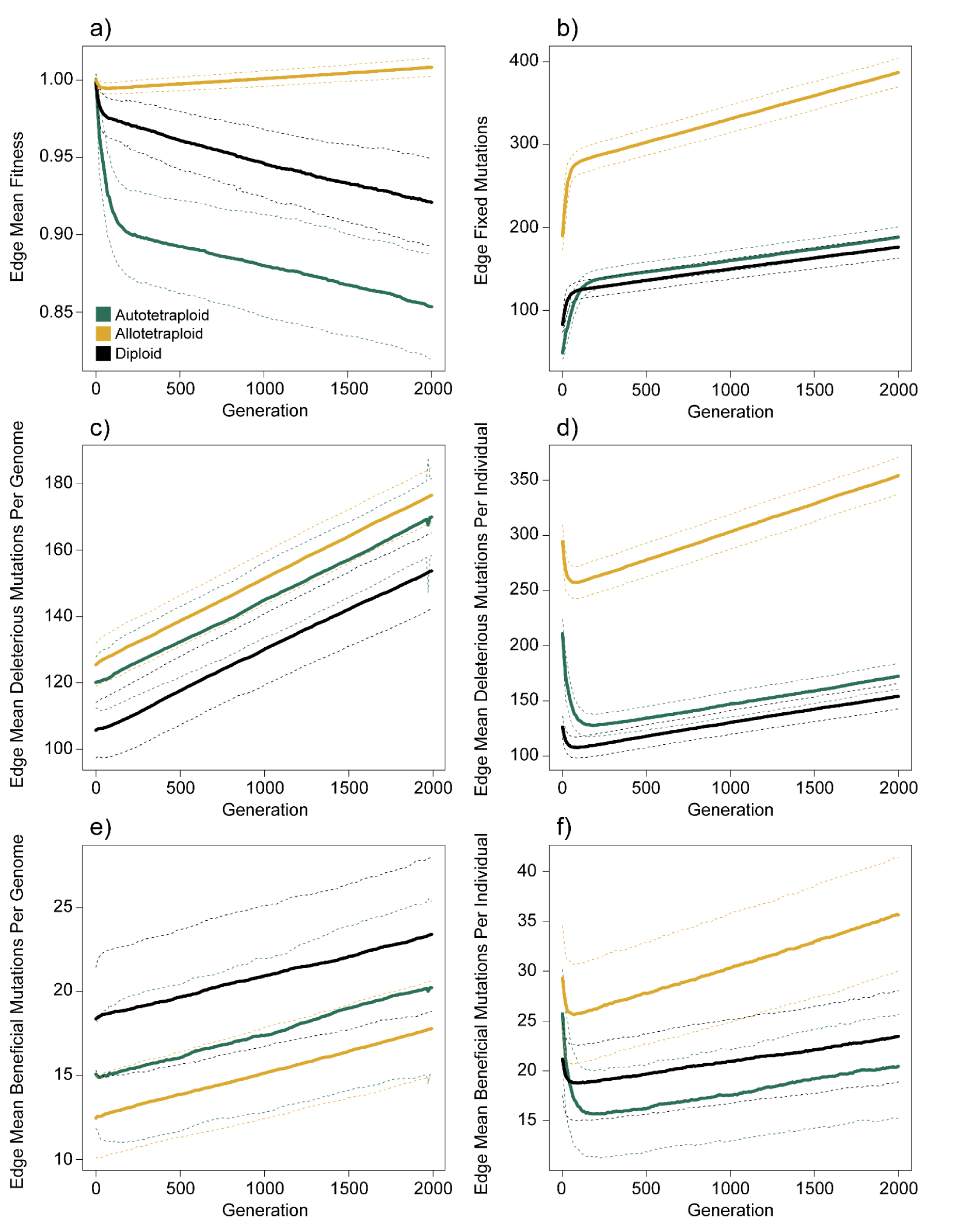
*h-s* relationship dominance model statistics. (a) Mean fitness (scaled) of the edge deme. (b) Number of fixed mutations in the edge deme. For allopolyploids, fixed mutations are those fixed within each subgenome. (c) Mean number of deleterious mutations per genome in the outermost deme (i.e. the edge). (d) Mean number of deleterious mutations per individual in the edge deme. (e) Mean number of beneficial mutations per genome in the outermost deme (i.e. the edge). (f) Mean number of beneficial mutations per individual in the edge deme. Lines show the average value of each statistic across 100 replicates, and dashed lines show ± 1 standard deviation.

**Figure 6.**
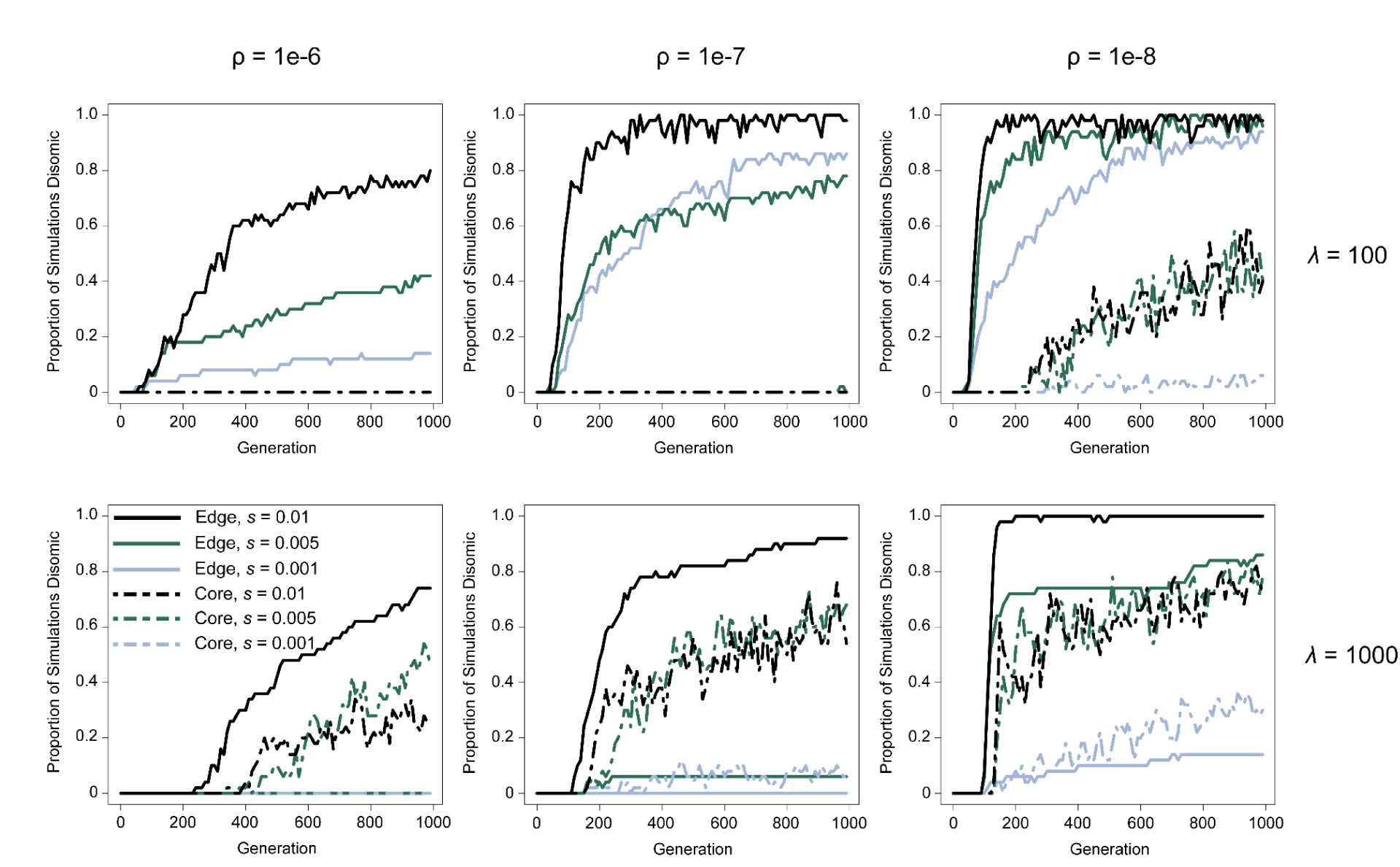
Diploidization of core and edge populations under the pairing efficiency model. Lines represent the proportion of simulations (out of 50 for each model) where diploidization evolved on the edge (solid lines) or core (dashed lines). Line color delineates the selective coefficient (*s*), and individual figures are separated by recombination rate (⍴; columns) and number of diploidization mutations (*λ* ; rows). *β* = 85 and δ = 1 for these simulations.

Although the incomplete recessiveness of the mutations in this model results in some fitness decline in allotetraploids, the magnitude of the fitness effects at 50% or lower frequency (the maximum frequency they can reach in allotetraploids) are miniscule and result in little difference between core and edge populations, and little fitness consequences generally.

### Cytogenetic diploidization is generally less likely on the expansion front unless disomy is restored by a single dominant mutation

Because recessive deleterious mutations fix rapidly in a spatially expanding autopolyploid with tetrasomic inheritance, we incorporated the ability to evolve disomy into our models to investigate if expansion increases the rate of cytogenetic diploidization in autopolyploids. Under a model in which disomy is restored by dominant mutations, the average rate at which populations diverge from random chromosomal associations (tetrasomy, diploidization index = 0) to complete subgenomic association (disomy, diploidization index = 1) is faster in core populations (Supp. Fig. 7). However, the average rate doesn’t accurately capture the stochastic dynamics of disomy evolution in expanding populations, where rapid fixation of diploidization alleles along the expansion front can lead to near instantaneous evolution of disomy along the edge. To investigate this, we considered the proportion of simulations where complete disomy (population diploidization index = 1) had evolved in core and edge populations. Here, the proportion of simulations where the entire population is disomic is frequently higher in edge populations, particularly in earlier generations and lower recombination rates when only 1 or 2 mutations are required to restore disomy (Fig. 5b). However, for the largest value of *λ* we examined, disomy is more likely to evolve in core rather than edgepopulations for any given ⍴ (with the exception of the case where ⍴ = 1×10^-8^ and *s*=0.01).

**Figure 5.**
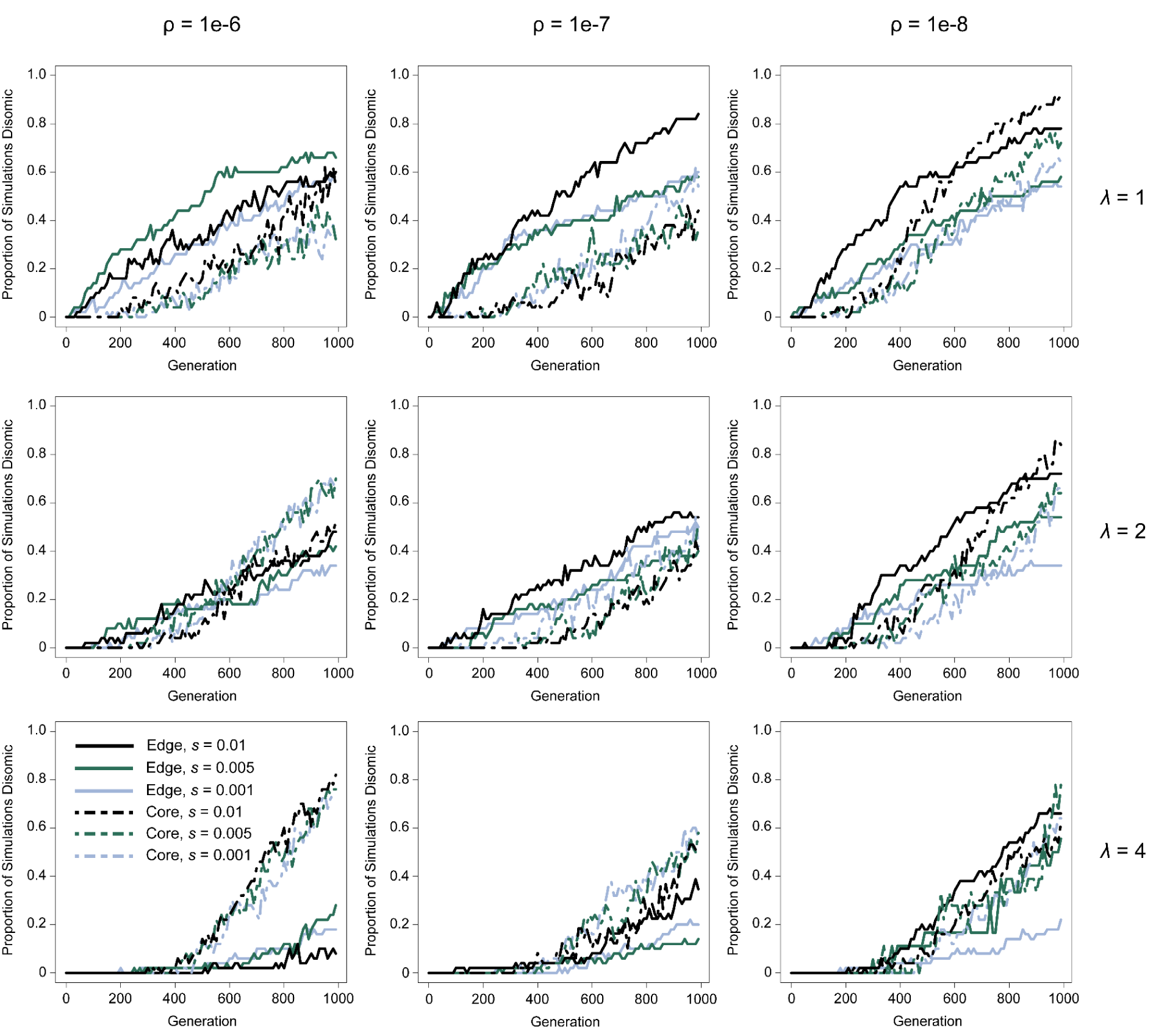
Diploidization of core and edge populations under the meiotic gene model. Lines represent the proportion of simulations (out of 50 for each model) where diploidization evolved on the edge (solid lines) or core (dashed lines). Line color delineates the selection coefficient (*s*), and individual figures are separated by recombination rate (⍴; columns) and the number of diploidization mutations required for full disomy (*λ*; rows).

We do not observe a clear relationship between disomy evolution and ⍴ for *λ* = 1 and 2. However at *λ* = 4, decreasing ⍴ below 1 recombination per chromosome per generation (1×10^-6^) seems to decrease the rate of disomy evolution in core populations, while the opposite appears to be true for the edge. There also does not appear to be a consistent impact of *s* on the disomy rate. However, a notable exception is observed when the recombination rate is low, and only 1 or 2 mutations are required for disomy: when *s*=0.01, the rate of disomy is elevated relative to the weaker selection coefficients along the edge for both ⍴=1×10^-7^ and ⍴=1×10^-8^, but only for ⍴=1×10^-8^ in core populations. In all, these results suggest that under a model where disomy is restored by mutations of large effect, disomy is less likely to evolve along the edge unless disomy can be restored with a single mutation, but somewhat equally as likely if the recombination rate is low.

An investigation into individual simulation replicates demonstrates the stochastic nature that expansion applies to the fixation of these mutations along the edge–in contrast with the more deterministic patterns found in the core (Supp. Fig. 8). For interior populations, the diploidization index fluctuates and may steadily evolve towards disomy. For edge populations however, changes to diploidization index happen quickly and are generally maintained in that and all subsequently colonized demes. Another interesting phenomenon individual simulation results show is that even if disomy doesn’t evolve on the edge more quickly, the stochastic dynamics occurring along the edge can have lasting effects long after those populations have established (e.g. maintained higher diploidization index in additional generations at *λ* = 2, 4; Supp. Fig. 8).

### Disomy evolves more rapidly on the expansion front under a wide range of conditions when pairing behavior is regulated by chromosomal similarity

While mutations of large effect have been previously discovered to restore disomy in several systems, evidence also suggests cytogenetic diploidization may proceed through chromosomal differentiation. Using a model adapted from Le Comber et al. (2010) where chromosomal similarity regulates pairing behavior with additional fitness costs, we find that diploidization is more likely along the expansion front under a large proportion of the parameter combinations investigated (Fig. 6, Supp. Figs. 9-12).

Although, *s*, ⍴, and the parameters of the pairing model (*β*, *δ*, and *λ*; see Methods) all had clear effects on the rate and nature of diploidization, *λ*, the number of loci determining chromosomal similarity, was the most significant determinant of whether disomy evolved more quickly in the core or edge (Fig. 6).

When *λ* = 100, disomy evolved more quickly on the edge for all other parameters investigated (Fig. 6, Supp. Figs. 9-10). Using the model parameters mimicking those from Le Comber et al. (2010), disomy evolved in a large proportion of the edge populations for all recombination rates, but only evolved in the core at the lowest recombination rate. Disomy was less likely to evolve for all parameters under the alternative models used, but when it did evolve, it did so more frequently in edge populations (Fig. 6). For the other more conservative models, disomy was less likely to evolve generally, and it is unclear whether disomy would have eventually evolved under some of these parameters because the length of simulations were limited by computation time. However when under the parameters where disomy did evolve under these models, it did so on the edge more quickly.

When *λ* = 1000, however, disomy was less likely to evolve along the edge except under higher values of *s* and lower values of ⍴ (Fig. 6). For higher values of *s* at higher ⍴, faster diploidization along the edge is often because, prior to diploidization, expansion has slowed due to reduced fitness along the range edge, and expansion only proceeds as fitter (and diploidized) individuals from the interior migrate to the expansion front. As well, for the more conservative *β* and δ models (i.e. models where *β* = 70), disomy was generally less likely to evolve, but usually did so more quickly in the core–again in contrast to the *λ* = 100 models (Supp. Figs. 11-12). However, at ⍴ = 1*e*-8, edge populations began diploidizing earlier for *s* = 0.01 and 0.005, although core populations would eventually catch up (outside of *β* = 70, δ = 0.2, and *s* = 0.01).

In contrast to the meiotic gene model, both the recombination rate and the selection coefficient of deleterious mutations have a clear effect in the pairing efficiency model for both core and edge populations, with the rate of diploidization generally increasing as the selection coefficient becomes more negative or the recombination rate decreases. Regarding recombination, as the recombination rate decreases the probability that diploidization loci are inherited together increases–increasing the number of differences between chromosomes causing disomy to evolve faster.

Regarding *s*, although our simulations of expansion load show a clear selective benefit of evolving disomy early along the edge to stop the fixation of deleterious recessive alleles (Fig. 3), the cause for diploidization in the core is less direct. However, as described in the Methods, the nature of this model is that there is a fitness cost if chromosomes are not either homogeneous in a unimodal distribution (tetrasomic) or in a bimodal distribution of two homogeneous groups (disomic). As such, because the model includes pairing-associated fitness costs (hereafter abbreviated as PAFC) there should be a fitness cost to the formation of chromosome pairs with high nucleotide diversity, thereby increasing the likelihood that a deleterious mutation will be found in the homozygous state. Indeed, in these simulations we observed a reduction in nucleotide diversity (Supp. Fig. 13a) and a correlated reduction in fitness (Supp. Fig. 13b) in the core. However this effect is delayed, with an initial increase in nucleotide diversity before the reduction because a certain level of differentiation between chromosomes must be achieved until there are fitness costs to pairing.

### Pairing associated fitness costs generate barriers to gene flow between disomic edge demes and tetrasomic interior demes

We further explored the dynamics of the pairing efficiency model by investigating models without PAFCs at varying selection coefficients including complete neutrality. Although we explored these dynamics for *λ* = 1000 and 100, all results were qualitatively similar and for brevity *λ* = 100 results are presented below. Interestingly, although the PAFCs are necessary for the evolution of disomy in core populations (Supp Figs. 14-15), PAFCs actually decrease the rate at which disomy evolves along the expansion front (Supp. Fig. 14). For all selection coefficients investigated, both the diploidization index and the proportion of simulations with restored disomy are higher along the edge when PAFCs are removed (although this state is transient, see below). In core populations, however, the diploidization index increases initially before returning to a lower baseline (Supp. Fig. 14b). Even with model parameters where disomy evolved in many core populations by 1000 generations (*β* = 85, δ = 1, *s* = -0.005, and ⍴ = 1×10^-8^), the diploidization index remained low in non-PAFC simulations (Supp. Fig. 15a) as the lack of fitness constraints results in pairing efficiency immediately falling towards 0 (Supp. Fig. 15b).

In contrast to core populations, pairing efficiency along the range edge is maintained near 1 throughout expansion with occasional sharp drops due to drift that are quickly recovered (Supp. Fig. 16). These drops generally coincide with an increase in the diploidization index, and occasionally are followed by the restoration of disomy. It is not clear why some but not all drops in pairing efficiency in tetrasomic populations result in the restoration of disomy. However, following the evolution of disomy, pairing efficiency is generally more stable around 1 along the edge, indicating minimal losses due to gamete failure once disomy is restored.

When disomy evolves in edge populations, this state is often maintained at that and any additionally colonized demes, indicating a fixation of this state on one side of the range edge (Supp. Fig. 17-18). Additionally, there is little spread of the disomic state to interior populations. When disomy evolves in interior populations, the spread is generally slow. One cause for this phenomenon may additionally be due to the PAFCs limiting migration, as F2 generations of offspring from tetrasomic and disomic parents may incur a fitness cost due to low pairing efficiency. If this is the case, we should expect a higher *F_ST_*between the populations along the tetrasomy-disomy border compared to other population comparisons. Indeed, we observe elevated *F_ST_* along that border in simulations with PAFCs (Supp. Fig. 19a), but no such elevated *F_ST_* when PAFCs are not included (Supp. Fig. 19b). Interestingly, we also observe higher *F_ST_* between adjacent demes that are disomic as opposed to between adjacent demes that are tetrasomic (Supp. Fig. 19a) indicating additional downstream effects once disomy evolves. When PAFCs are removed from the model, *F_ST_* between adjacent demes is minimal, including along the tetrasomy-disomy border (Supp. Fig. 19b). One result of this increased effective rate of gene flow is that, without PAFCs, disomy along the edge is transient, although it is also possible that reversion to tetrasomy is expected without PAFCs even in an isolated population–a scenario untested here.

## DISCUSSION

### The genetic consequences of range expansion in polyploids

The results of this study have several implications for our understanding of polyploid evolution and establishment. Perhaps most immediately is the relationship between inheritance and expansion in polyploids, introducing an additional layer of complexity to the longstanding hypothesis that polyploidy confers an advantage to colonizing novel and deglaciated habitats. More specifically, outside of an unlikely model of additive dominance, the masking advantage provided by whole genome duplication when mutations are partially or fully recessive results in an accumulation of deleterious mutations that in turn leads to a protracted drop in the fitness of polyploids with tetrasomic inheritance during expansion relative to their diploid counterparts. Indeed, recent evidence in Coho salmon (*Oncorhynchus kisutch*) showed residual tetraploid regions of their genome that had not yet re-diploidized harbored a greater mutational load than the diploidized regions (Rougemont et al. 2023)

Abundant evidence has demonstrated a link between polyploidy and previously glaciated areas (Brochmann et al. 2004; Paun et al. 2006; Novikova et al. 2018; Sutherland and Galloway 2018; Rice et al. 2019; David 2022; Booker et al. 2023). While support for this observation has many adaptive explanations–such as fixed heterozygosity (Stebbins 1985; Brochmann et al. 2004), increased adaptability of gene regulatory networks (Freeling and Thomas 2006; Hegarty and Hiscock 2007; Fusco et al. 2010; Yao et al. 2019; Ebadi et al. 2023), or the relaxed constraint on gene duplicates enabling more rapid adaptation (Lynch and Conery 2000; Lynch and Force 2000; Gout and Lynch 2015)–these arguments too rely on some level of subgenomic differentiation, and therefore disomy, to maintain their validity. As such, the results herein suggest polyploids with tetrasomic inheritance should be uniquely disadvantaged in colonizing new areas as polyploidy confers significant consequences as a result of expansion load–not to mention additional disadvantages due to meiotic errors as a result of polyploidy (Mayer and Aguilera 1990; Comai et al. 2000; Ramsey and Schemske 2002; Morgan et al. 2020). However, because the work here investigates the consequences of expansion alone, more work extending the scenarios examined here to include other potential advantages of polyploidy would be required to determine whether or not the adaptive benefits to polyploidy may in some cases offset the disadvantages that we show here.

One potential way to offset the disadvantages inherent to autopolyploids expanding their range is through interploid hybridization. Despite the fitness consequences of hybridizing across ploidies, numerous studies have demonstrated pervasive gene flow between autopolyploids and their lower-ploidy relatives when their ranges overlap (Arnold et al. 2016; Marburger et al. 2019; Monnahan et al. 2019; Bogart et al. 2020; Novikova et al. 2020; Shastry et al. 2021; Booker et al. 2022). Although hybridization across ploidies during formation is often considered a way for neo-polyploids to overcome the massive population bottleneck during formation (Stebbins 1980), admixture following their establishment should be generally disfavored as mixed-ploidy offspring suffer extensive fitness consequences (Ramsey and Schemske 1998)–though this may be mitigated in higher-level ploidies (Peskoller et al. 2021; Sutherland and Galloway 2021). However, hybridization to relieve the deleterious load accumulated through range expansion may help to explain why this phenomenon is so pervasive, and why the evolutionary histories of autopolyploids can be so reticulate (e.g. Booker et al. 2022). Importantly, interploid hybridization has also been documented in numerous allopolyploids (Ma et al. 2010; Sutherland and Galloway 2018; Kryvokhyzha et al. 2019b; Kryvokhyzha et al. 2019a; Zhao et al. 2019). While disomic neo-allopolyploids should be shielded from the effects of expansion, some level of tetrasomy (as is often observed in allopolyploids, see Li et al. 2021) or fractionation should make some degree of post-expansion interploid hybridization more likely.

While autopolyploids may be uniquely disadvantaged in expanding their range under realistic models of dominance, the results of this study show a markedly different pattern in allopolyploids. Allopolyploids may be at a disadvantage under an additive model, though this is largely due to fitness consequences incurred in isolation. However the restriction of recombination across subgenomes prevents the fixation of alleles as a result of expansion (unless the same mutation occurs on both subgenomes), therefore providing a benefit if deleterious mutations are fully or partially recessive. Importantly, the model investigated here is of a neo-allopolyploid where no fractionation or genic diploidization, wherein gene duplicates are gradually lost or silenced leading to dynamics more and more similar to diploids, has occurred. In reality, allopolyploids exist somewhere on the continuum between the neo-allopolyploids investigated here and a functionally diploid paleo-polyploid. Absent additional genomic alterations as a result of polyploidization, anywhere outside of the functionally diploid paleo-polyploid extreme allopolyploids should suffer fewer consequences of expansion load relative to diploids. However, because the process of polyploidization itself results in numerous additional genomic consequences, there are likely additional factors affecting an allopolyploid’s response to range expansion that remain unexplored here. Nevertheless, results from this study may provide some insight into these additional alterations: because disomic inheritance results in the accumulation of fixed deleterious mutations within each subgenome (Fig. 4e), these dynamics may have consequences for the process of fractionation (gene silencing, subfunctionalization, and loss) in later stages of polyploid evolution. Furthermore, because genes on one subgenome may have a high number of deleterious mutations, this may hinder the adaptability of allopolyploids through functional redundancy.

Although the present study focuses on the consequences of expansion for polyploids already formed, the work here and elsewhere provokes a reconsideration of the hypotheses purported by Stebbins (1984, 1985) and evidenced by Brochmann et al. (2004): that allopolyploid formation is promoted in the arctic as glacial cycles promote secondary contact which can fix heterozygosity in newly formed polyploids. Macpherson et al. (2022) showed that range expansion promotes hybridization of species when they come in contact by providing immediate relief to the genetic load built up by expansion, and under some parameters the benefit provided by hybridization is enough to counteract the negative effects of Bateson-Dobzhansky-Muller incompatibilities.

Allopolyploid formation rather than diploid hybridization provides these same benefits, while also introducing strong reproductive isolation. As demonstrated here, should these newly formed allopolyploids have largely disomic inheritance, their formation here would not only provide immediate relief to the already fixed load incurred by the range-expanding diploids, but they would also be protected from the build up of additional load as expansion continues. Although the establishment and persistence of a new polyploid under these circumstances is still a strong evolutionary hurdle to overcome, the benefits relative to diploids regarding expansion load should make this more likely.

The terms we use here as autopolyploid and allopolyploid provide shorthand for tetrasomically and disomically inheriting polyploids. The definition of these terms are not exact, often referring to the nature of their formation or behavior in meiosis. In reality, nearly half of all identified allopolyploids have some level of multivalent formation (Li et al. 2021) and therefore necessarily deviate from the disomic inheritance modeled in allopolyploids here. As such, the particular consequences of expansion prescribed to autotetraploids are generally applicable across polyploids to the degree which their inheritance allows deleterious mutations to fix across all chromosomal copies. Similarly, the benefits described here to allotetraploids translate to the extent in which recombination across homoeologs is restricted, and many tetraploids may experience dynamics somewhere in between those of the fully tetrasomic and disomic tetraploids examined here. Additionally, although this work focuses on tetraploids, the masking potential of polyploidy–and the consequences of that masking when those species expand their range–should be generally applicable to polyploids of higher ploidy as well.

### Cytogenetic diploidization of range-expanding polyploids

We modeled the dynamics that regulate the inheritance pattern of polyploid meiosis during a range expansion and observed several interesting behaviors related to both the nature of diploidization and the fitness consequences of expansion. Regardless of the method by which cytogenetic diploidization proceeds, results from this study show that expansion can increase the rate at which disomy is restored in a polyploid under a wide range of parameters. Although the models of diploidization explored here are fairly simple, they represent two empirically identified modes of cytogenetic diploidization in nature. The better-understood of these two models is the meiotic gene model, which incorporates dominant mutations that alter the meiotic machinery itself. Among these *Ph1*, a gene identified in grasses that when present suppresses pairing of homoeologs and facilitates pairing of homologs by affecting the interactions between microtubules and kinases (Vega and Feldman 1998), has been the most extensively studied (Riley and Chapman 1958; Riley 1960; Sears 1976; Sánchez-Morán et al. 2001; Feldman and Levy 2012; Li et al. 2021). A similar gene that experiences mutations of large effect which regulate pairing behavior is *PrBn* in *Brassica napus* (Jenczewski et al. 2003), and an additional QTL of large effect has also been found in *Arabidopsis suecica* (Henry et al. 2014). However, other identified pathways to the genetic control of meiosis have also been observed, such as *MHS4* copy number regulation that may be preserved in angiosperms generally (Gonzalo et al. 2019) and a complex landscape of epistatic interactions in *Arabidopsis arenosa* and *A. thaliana* neo-autopolyploids (Morgan et al. 2020; Morgan et al. 2022; Gonzalo et al. 2023).

Under the meiotic gene model, the evolution of disomy is largely stochastic and minimally dependent on the fitness consequences of expansion (Fig. 5b,c). Instead, the rate of evolution of disomy is determined by the number of mutations required to restore disomy, and the chance that any already-segregating mutations fix is higher on the expansion edge. As well, even if disomy is not completely restored along the edge when *λ* > 1, any mutations that influence inheritance and that happened to fix along the edge may persist as the range continues to expand and the former edge populations become interior populations (Supp. Fig. 8). Thus, the stochastic fixation of disomy-restoring mutations along the range edge may increase the rate at which disomy evolves in the species across a broad portion of its range.

While our best-known examples of meiotic regulation in polyploids come from genetically controlled mechanisms such as those described above, it is not necessarily the case that this is the predominant mechanism of cytogenetic diploidization as identifying the relative contribution of chromosomal differentiation to pairing behavior (i.e. the pairing efficiency model) is inherently more difficult. Still, competition experiments have shown that pairing preferences are linked to some aspect of sequence divergence in certain systems (Bomblies 2022). Indeed, both large structural changes, such as inversions (Parra-Nunez et al. 2018), and the more uniform accumulation of sequence divergence along the chromosome (Braz et al. 2021) have been shown to result in preferential pairing.

We found the influence of expansion on the evolution of disomy through the pairing efficiency model to be patently different from that of the meiotic gene model. Under this model, cytogenetic diploidization is in many ways more predictable and the rate of disomy restoration is linked to the selection coefficient in both core and edge populations (Fig. 6b,c). Although this was not the original purpose of the present study, the result that adding deleterious recessive mutations to the model of Le Comber et al. (2010) increases the speed in which disomy is restored lends additional credence to the validity of this model as a pathway to diploidization. Furthermore, this result also makes this model less reliant on subfunctionalization and neofunctionalization as a necessary accelerator of diploidization.

Deleterious recessive mutations influence the rate of disomy restoration because during the tetrasomic phase of a population there is a fitness consequence (PAFCs) to heterozygosity, and selection acts to increase homozygosity generally–increasing the probability recessive mutations will be found in the homozygous state as well. While this acts to reduce fitness due to deleterious variation generally, it also introduces a selective pressure to transition to disomy, where high levels of nucleotide diversity are no longer disadvantageous. This results in fitness losses in the core during the transition to disomy, whereas along the edge we observe a less dramatic fitness loss because the transition to disomy also works to counteract the buildup of recessive mutations from expansion (Supp. Fig. 13b).

Although the pairing efficiency model is largely deterministic in nature, it is the outcome of stochastic events that govern that nature. This is first evidenced by the increasing rate of diploidization as the recombination rate decreases (Fig. 6). Here, lower recombination rates increase the haplotype block length, thereby increasing the number of jointly inherited pairing mutations. This maximizes the distance crossed on the pairing efficiency curve in a single generation making diploidization more likely. Similarly, the more impact individual pairing mutations have (i.e. lower *λ*), the more impact the chance fixation of those mutations on the expansion front can have. Translated to real populations, this result means that mutations of large effect (e.g. large duplications, inversions, translocations) or the buildup of genetic variation in low-recombining regions will tend to increase the probability that expansion results in a faster transition to disomy, whereas SNPs and other small mutations in higher recombining areas are less likely to do so. Interestingly, this pattern is reversed in the core, where a more gradual buildup of smaller-effect mutations makes diploidization more likely.

Importantly, the empirical examples above discuss pairing behavior as it relates to both chiasma formation (e.g. bivalents or multivalents) and inheritance. While the pairing efficiency model indirectly includes some consequences of multivalent formation (PAFCs), we do not explicitly model multivalent formation as part of these models. In large part this is to simplify the models and identify the influence of range expansion on disomy restoration as a potential conduit of diploidization. Evolving disomy can be viewed as an extreme alteration to the recombination landscape of an organism, and additional work that might identify other routes of recombination modification such as modeling multivalent formation, more realistic recombination maps, and dynamic recombination rates and landscapes as chromosomes diverge may provide additional routes through which an expansion influences diploidization of polyploids.

### Conclusions

The present work was conducted to examine aspects of polyploid evolution and establishment under two processes all polyploids proceed through: range expansion and diploidization. To do so, we have introduced more realistic models that allowed us to observe behaviors that do not follow additively from these extensions in complexity. In one case the behavior renders an otherwise advantageous element disadvantageous: in autopolyploids, the duplicated genome provides a fitness benefit to an isolated population through masking recessive mutations, but this same mechanism in turn results in severe fitness consequences as deleterious mutations accumulate along the range edge as the population expands. In other instances we made observations that were wholly unexpected. For example, if individual mutations have large effects on pairing behavior, disomy evolves faster along the expansion front but slower in core populations, with the opposite being true for mutations of relatively smaller effect. Additionally, by modeling cytogenetic diploidization spatially, we found that asynchronous evolution of disomy across the landscape can create barriers to gene flow–a behavior that could result in future speciation. Spatial dynamics play an important role in the evolutionary trajectory of all species, but the conditions that regulate polyploid formation and establishment suggest an elevated importance of this role in polyploid evolution. We anticipate that future work incorporating these dynamics into population genetic models of polyploids will continue to help answer remaining questions and paradoxes regarding polyploid evolution.

## ACKNOWLEDGEMENTS

We would first like to thank Ben Haller and Peter Ralph for providing prompt and necessary feedback while developing the polyploid models for these simulations. We would also like to thank the attendants of the 2023 Polyploidy Tree of Life Conference for their helpful comments and feedback of this work as it was ongoing, Tyler Kent for providing valuable feedback on this manuscript and discussion as the project was ongoing, Justin Conover for insightful comments and discussion on the initial preprint of this work, and the members of the Schrider lab for their input throughout this project. This work was supported by NIH grants HG010774 and GM138286.

## DATA ACCESSIBILITY

All scripts used for this study can be found in the github repository https://github.com/wbooker/polyploid_expansion_load

## Supplemental Information

**Supplemental Figure 1.**
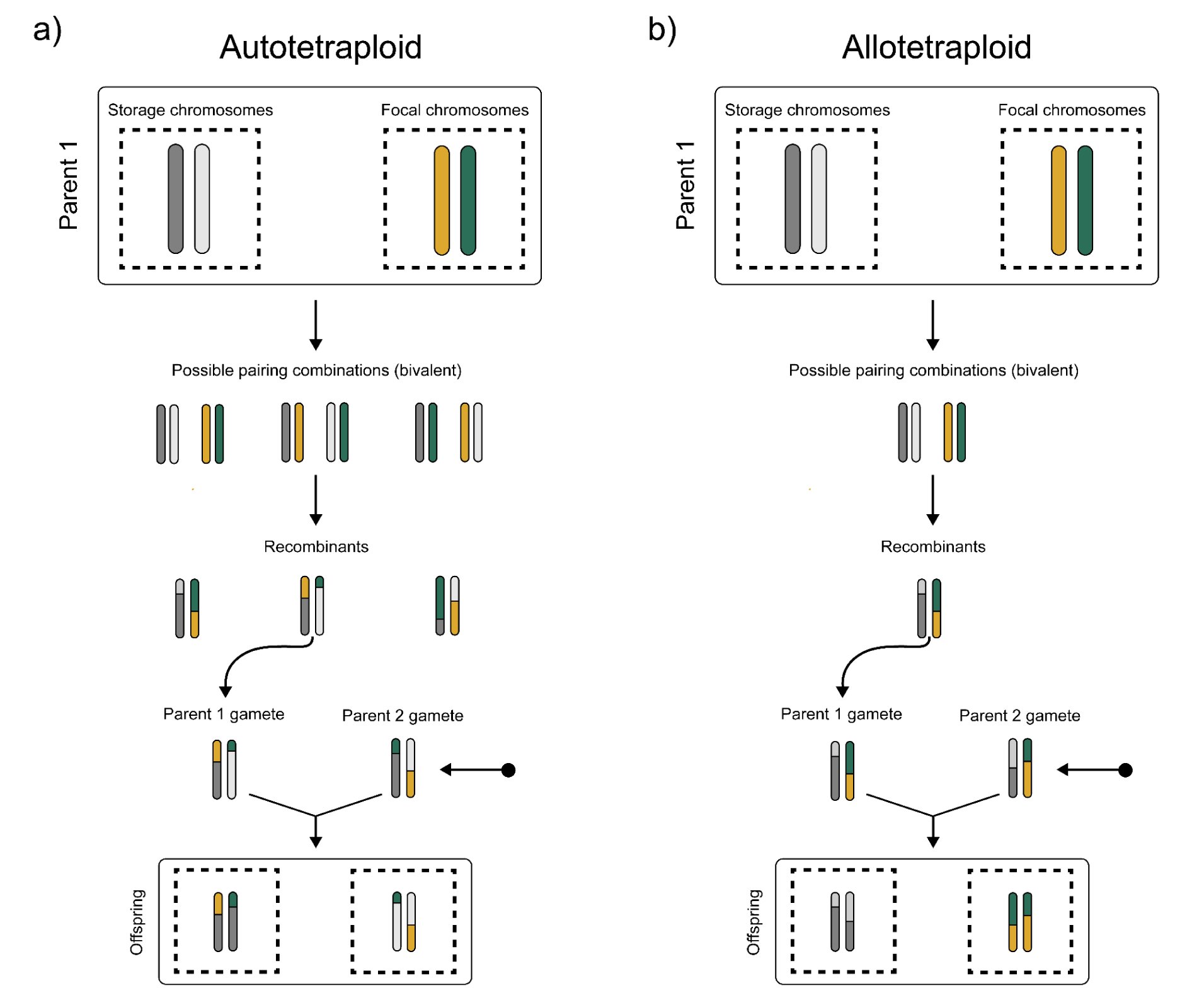
Example diagram of reproduction in (a) autotetraploid and (b) allotetraploid as modeled in our SLiM simulations. In autotetraploids, there are three possible pairing combinations of the four chromosomes given the bivalent pairing modeled here. A combination is chosen at random, and recombinant chromosomes are combined into the offspring without regard as to which recombinants are placed in the new storage and focal populations. In allotetraploids, there is only one possible pairing, and recombinant chromosomes maintain their affinity to their original population so as to prevent homoeologous exchange.

**Supplemental Figure 2.**
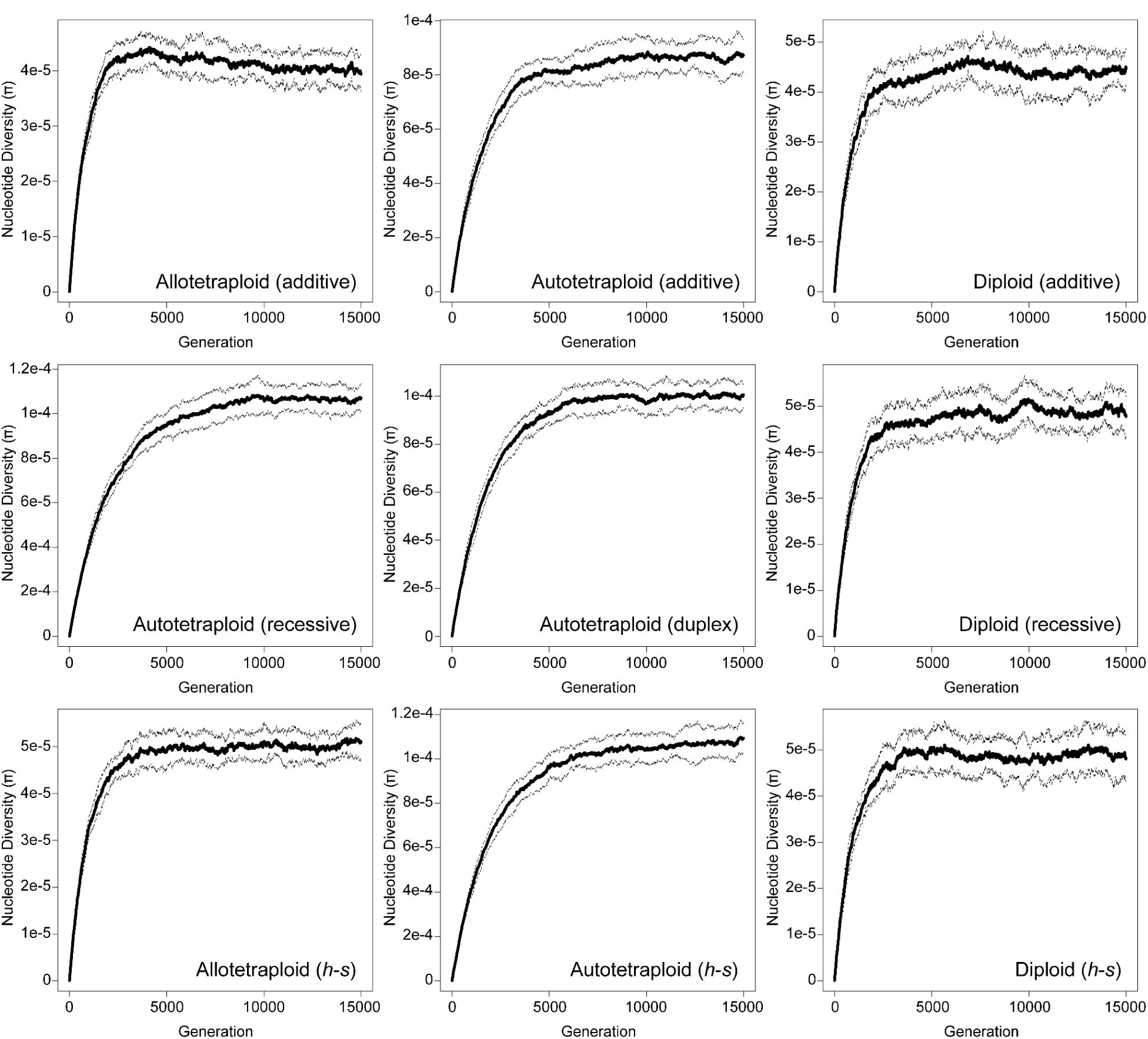
Nucleotide diversity (*π*) in the core as a function of the number of burn-in generations for all expansion models. Solid lines represent the average nucleotide diversity across 20 generations, with dashed lines representing ± 1 standard deviation.

**Supplemental Figure 3.**
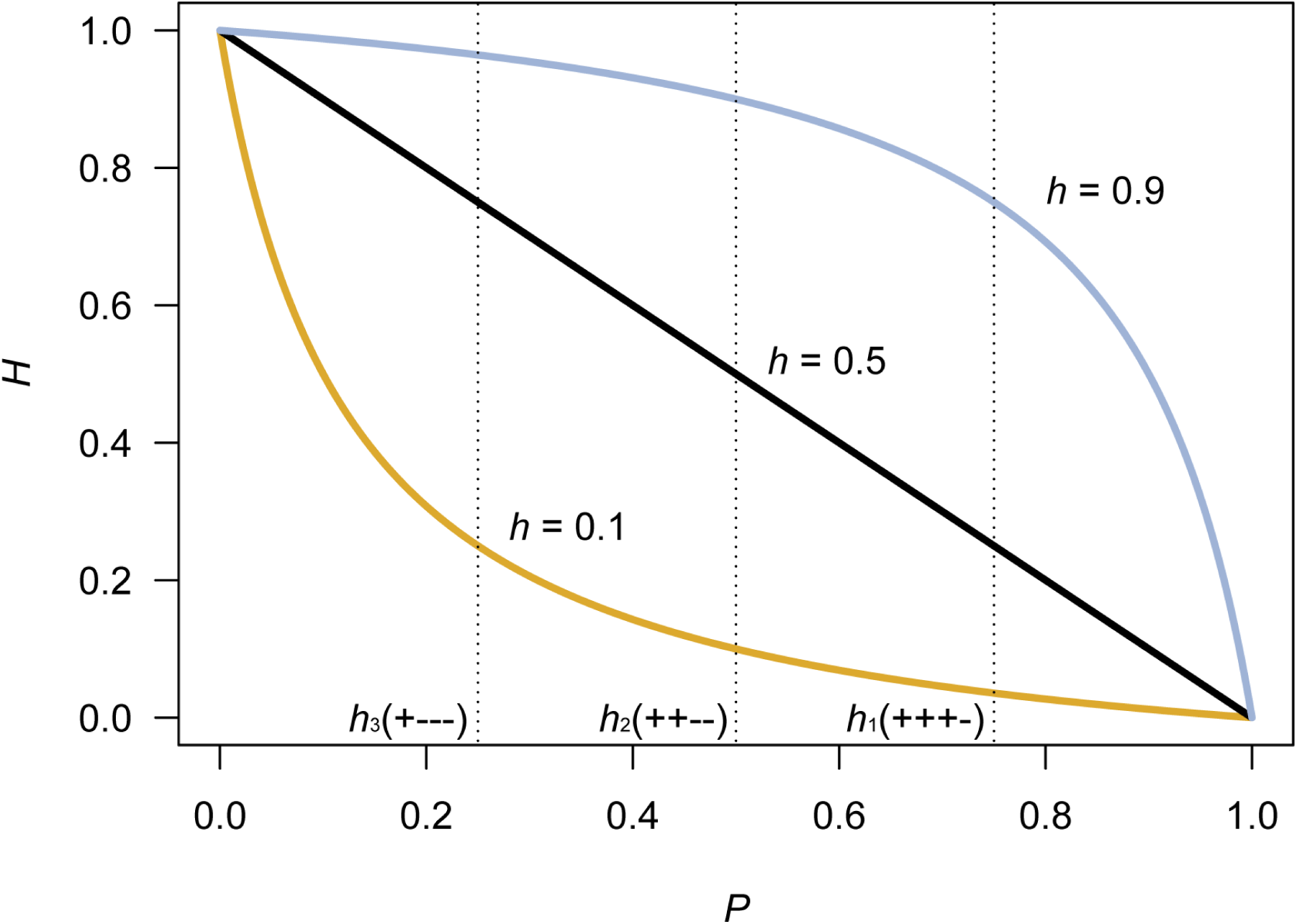
The relationship between the continuous dominance coefficient (*H*) given the frequency of wild-type alleles (*P*) following the Kacser and Burns (1981) model of flux. Curves depict the values of *H* for example diploid dominance coefficients for partially recessive (*h* = 0.1), additive (*h* = 0.5), and partially dominant (*h* = 0.9) mutations. Dashed lines show sampled *H* values for the tetraploid *h* vector as in Table 1, given the number of wild-type (+) and mutant (-) alleles.

**Supplemental Figure 4.**
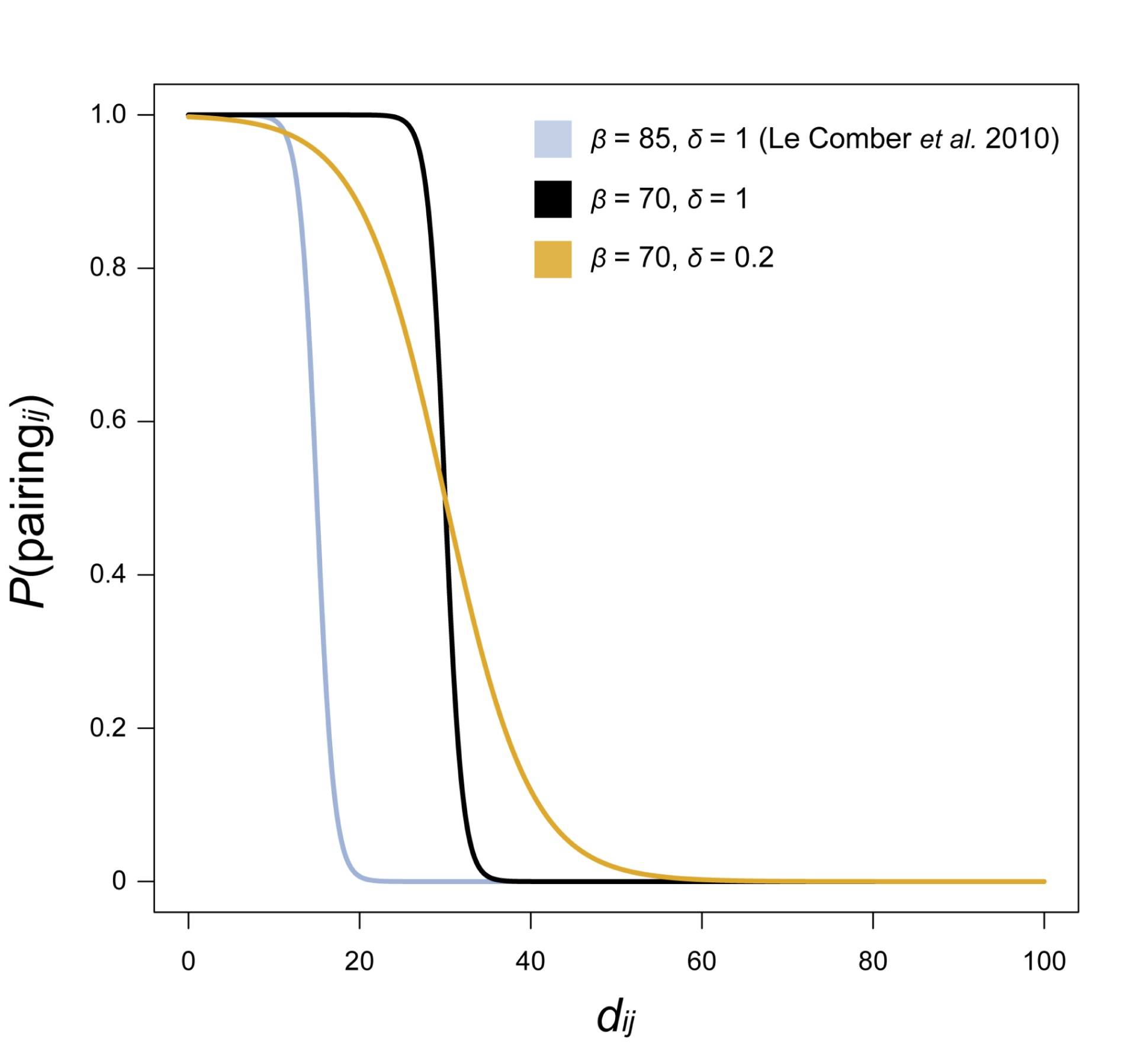
Probability of two chromosomes *i* and *j* pairing (i.e. pairing efficiency) based on their percent divergence (*d_ij_*) as defined in Eq. 10 for three parameter combinations of the inflection parameter *β* and slope δ.

**Supplemental Figure 5.**
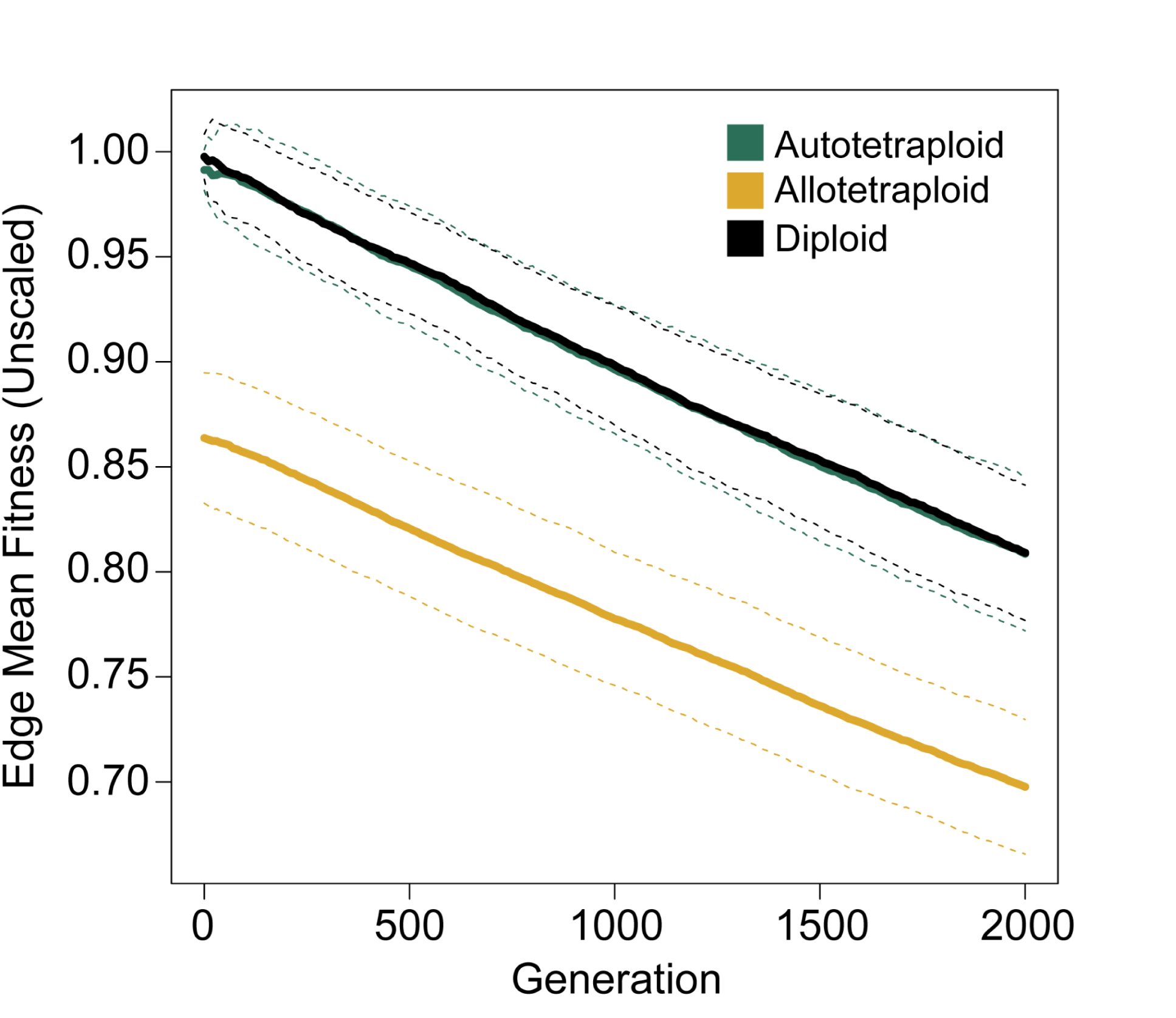
Unscaled mean fitness of the edge deme for additive models with no beneficial mutations.

**Supplemental Figure 6.**
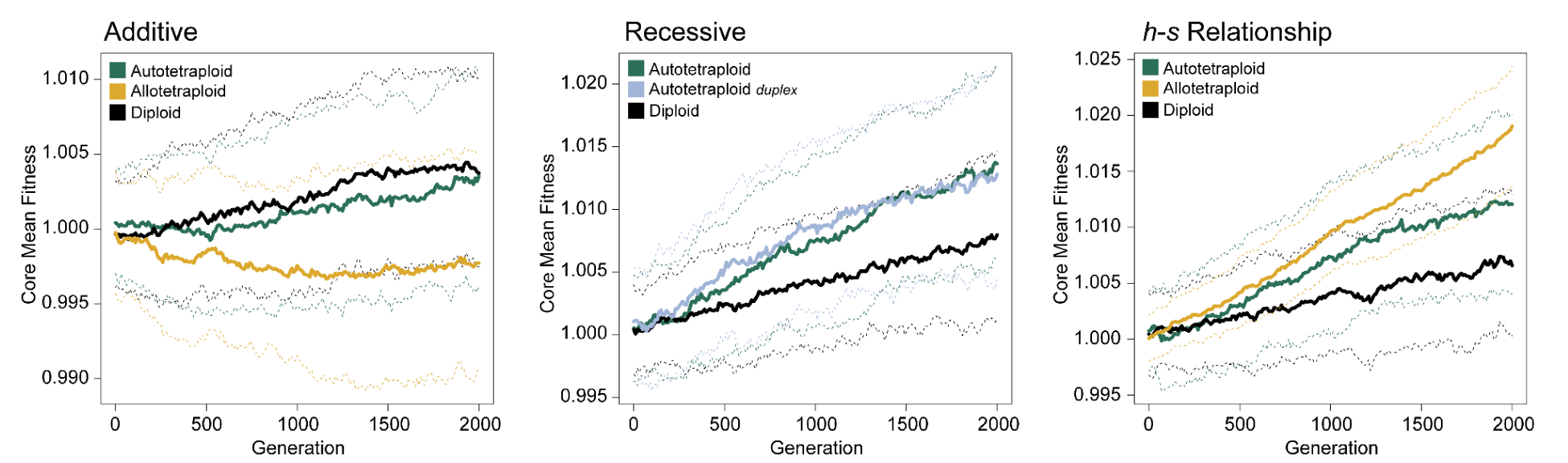
Mean fitness (scaled) in the core for each of the dominance models investigated. All means represent 100 simulations, dashed lines show ± 1 standard deviation.

**Supplemental Figure 7.**
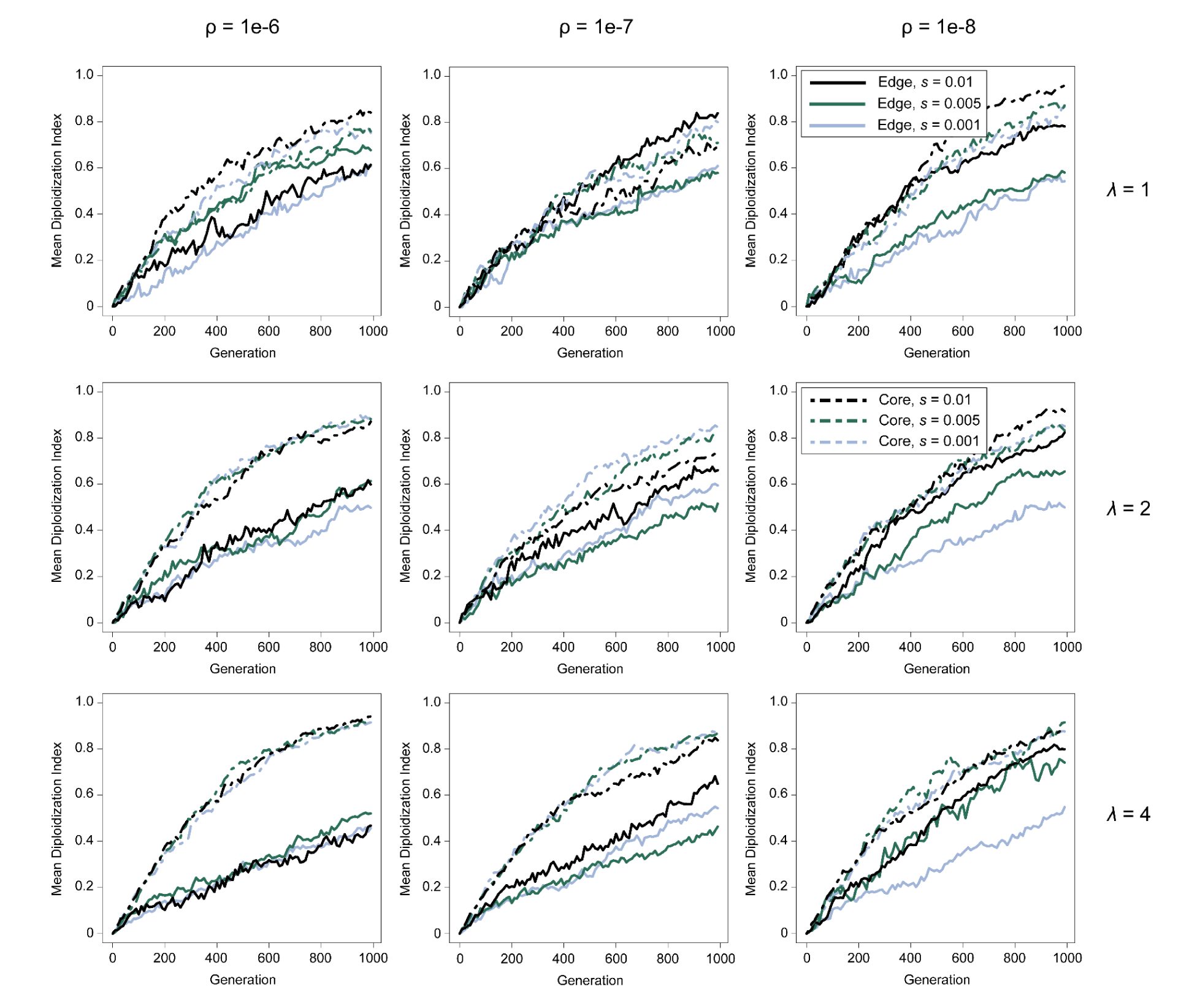
Mean diploidization index of core (dashed lines) and edge (solid lines) populations under the meiotic gene model.

**Supplemental Figure 8.**
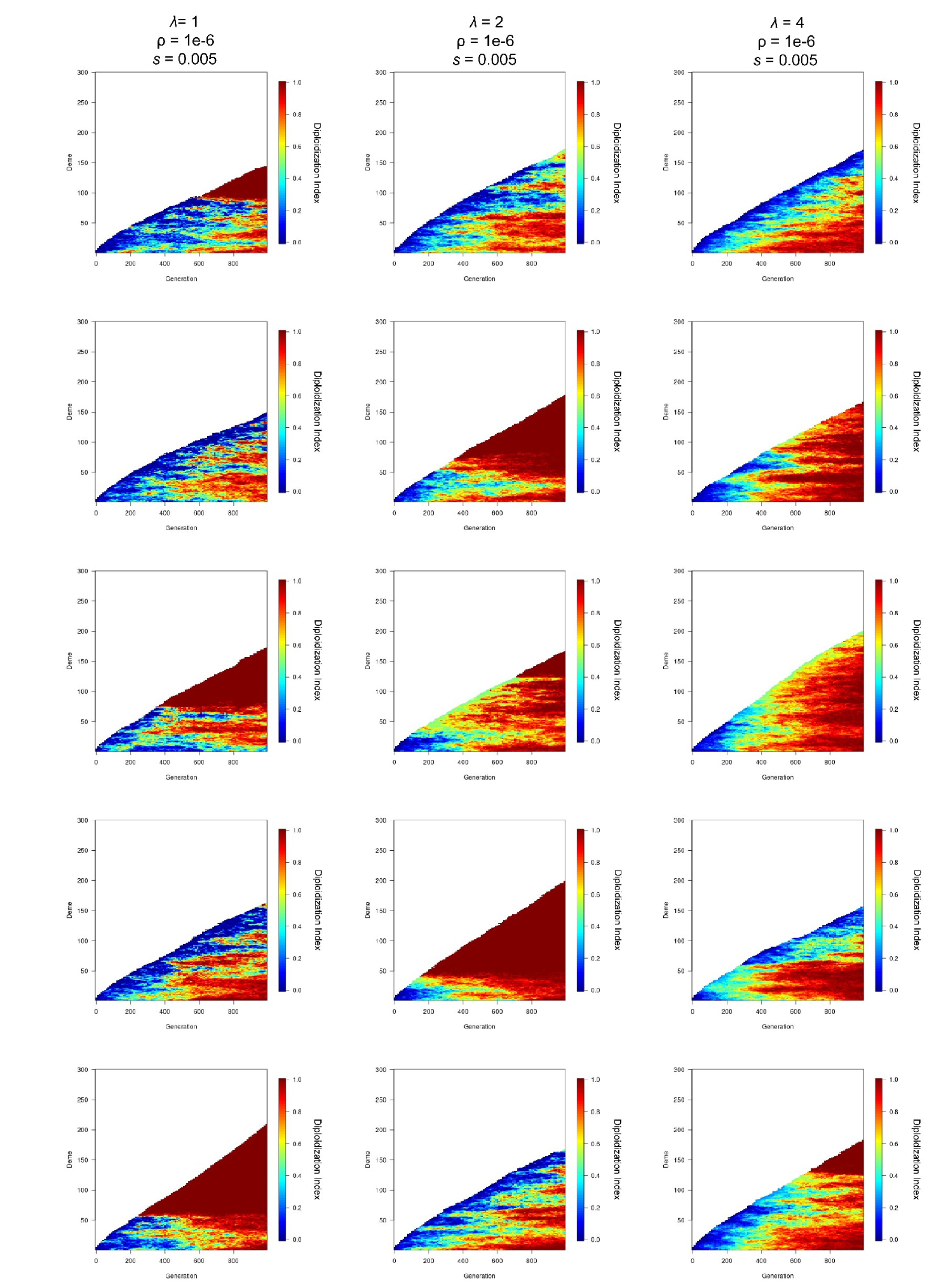
Diploidization index of example simulations under the meiotic gene diploidization model with 1-4 mutations required to restore disomy.

**Supplemental Figure 9.**
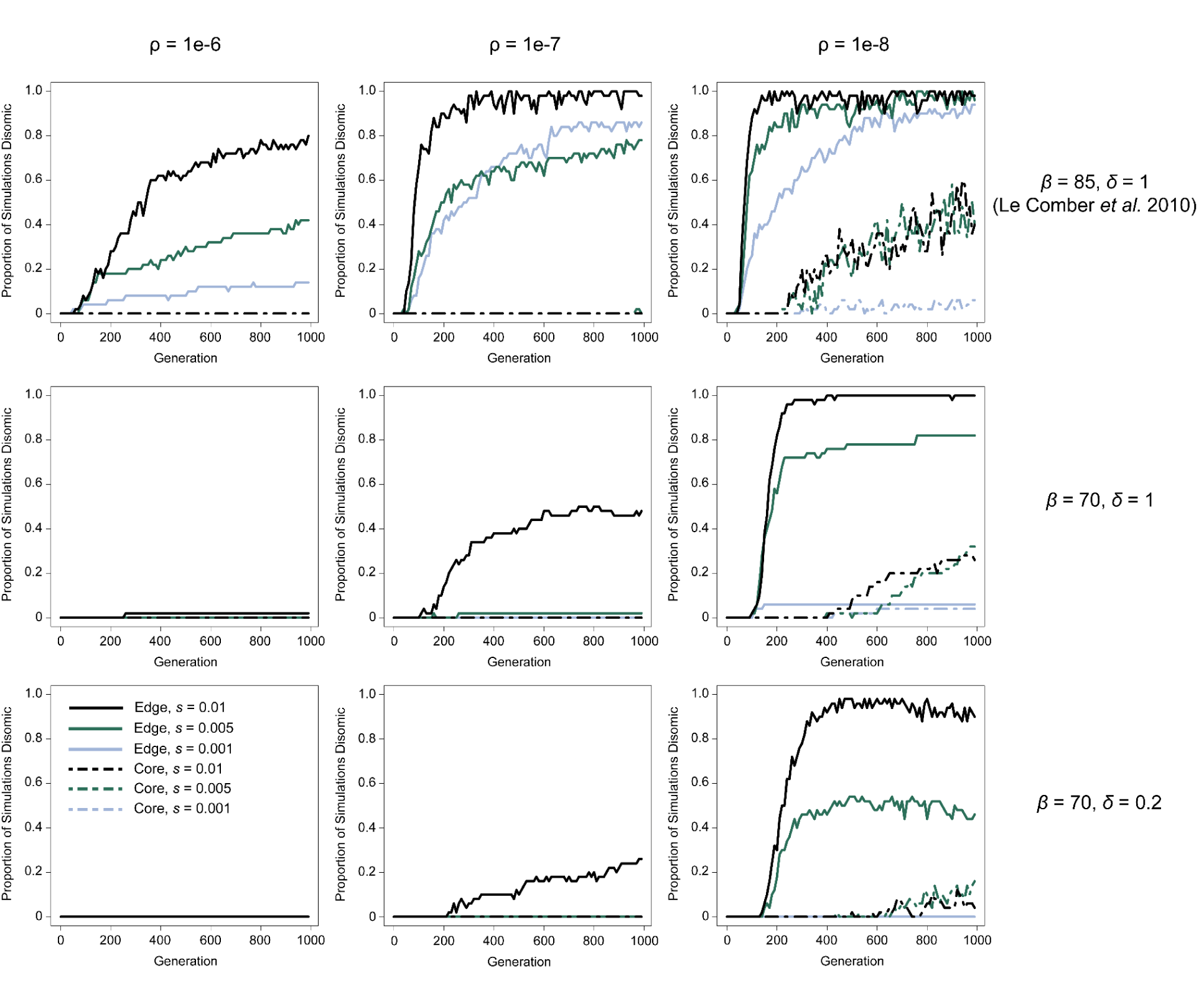
Diploidization of core and edge populations under the pairing efficiency model at *λ* = 100. Lines represent the proportion of simulations (out of 50 for each model) where diploidization evolved on the edge (solid lines) or core (dashed lines). Line color delineates the selective coefficient (*s*), and individual figures are separated by recombination rate (⍴; columns) and the parameter set for the model full disomy (*β* and δ; rows).

**Supplemental Figure 10.**
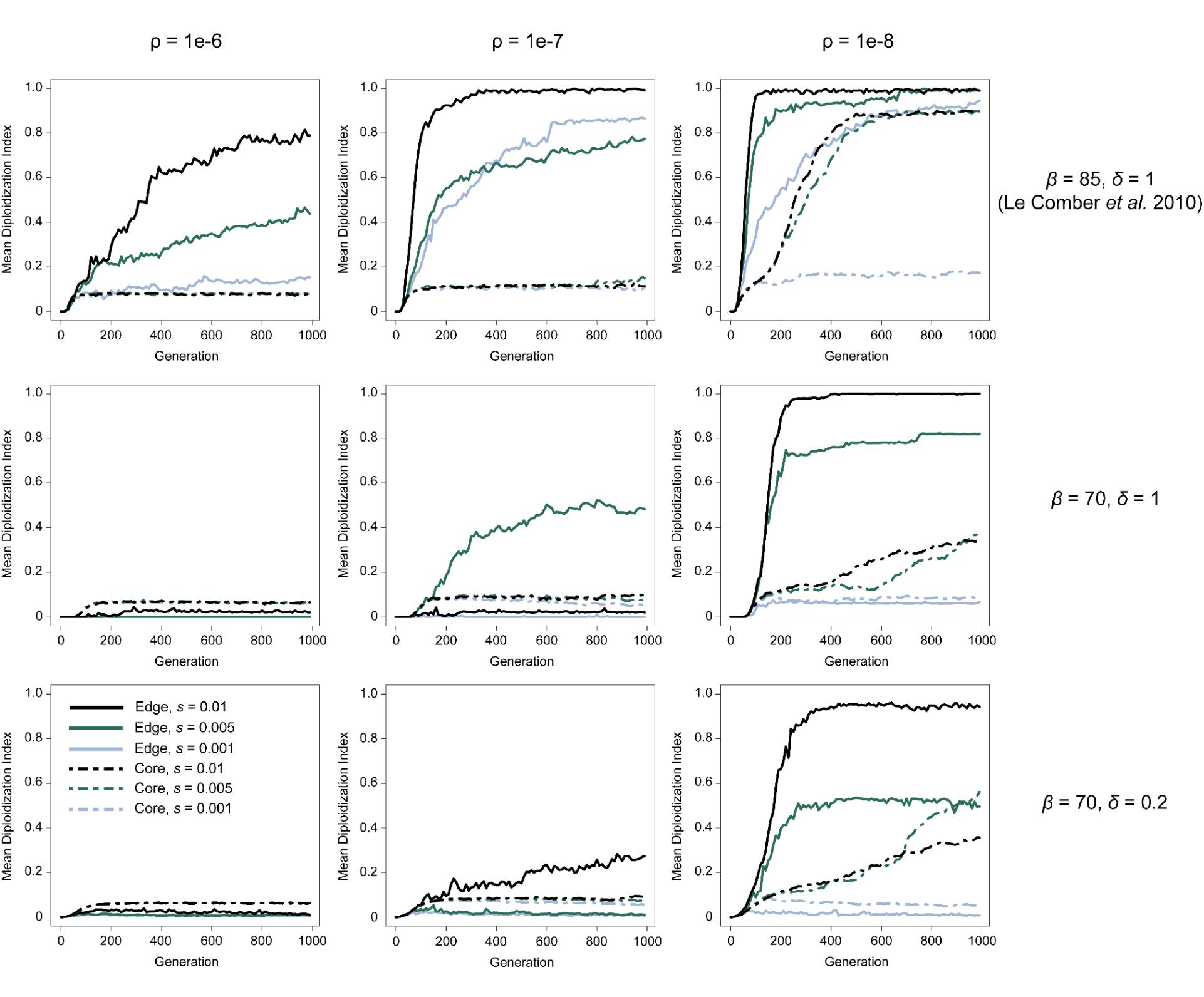
Diploidization of core and edge populations under the pairing efficiency model at *λ* = 100. Lines represent the mean diploidization index (out of 50 for each model) in the edge (solid lines) or core (dashed lines). Line color delineates the selective coefficient (*s*), and individual figures are separated by recombination rate (⍴; columns) and the parameter set for the model full disomy (*β* and δ; rows).

**Supplemental Figure 11.**
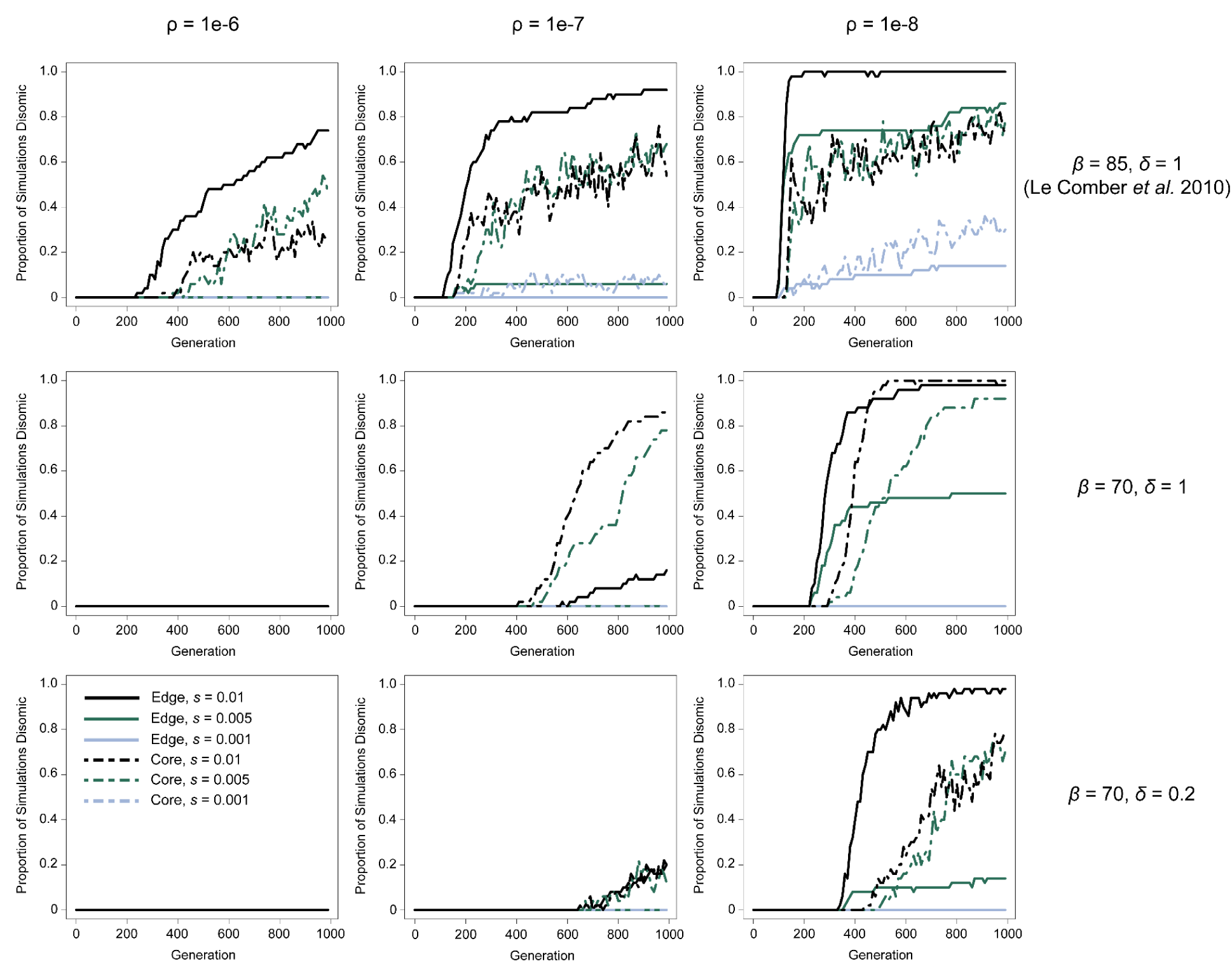
Diploidization of core and edge populations under the pairing efficiency model at *λ* = 1000. Lines represent the proportion of simulations (out of 50 for each model) where diploidization evolved on the edge (solid lines) or core (dashed lines). Line color delineates the selective coefficient (*s*), and individual figures are separated by recombination rate (⍴; columns) and the parameter set for the model full disomy (*β* and δ; rows).

**Supplemental Figure 12.**
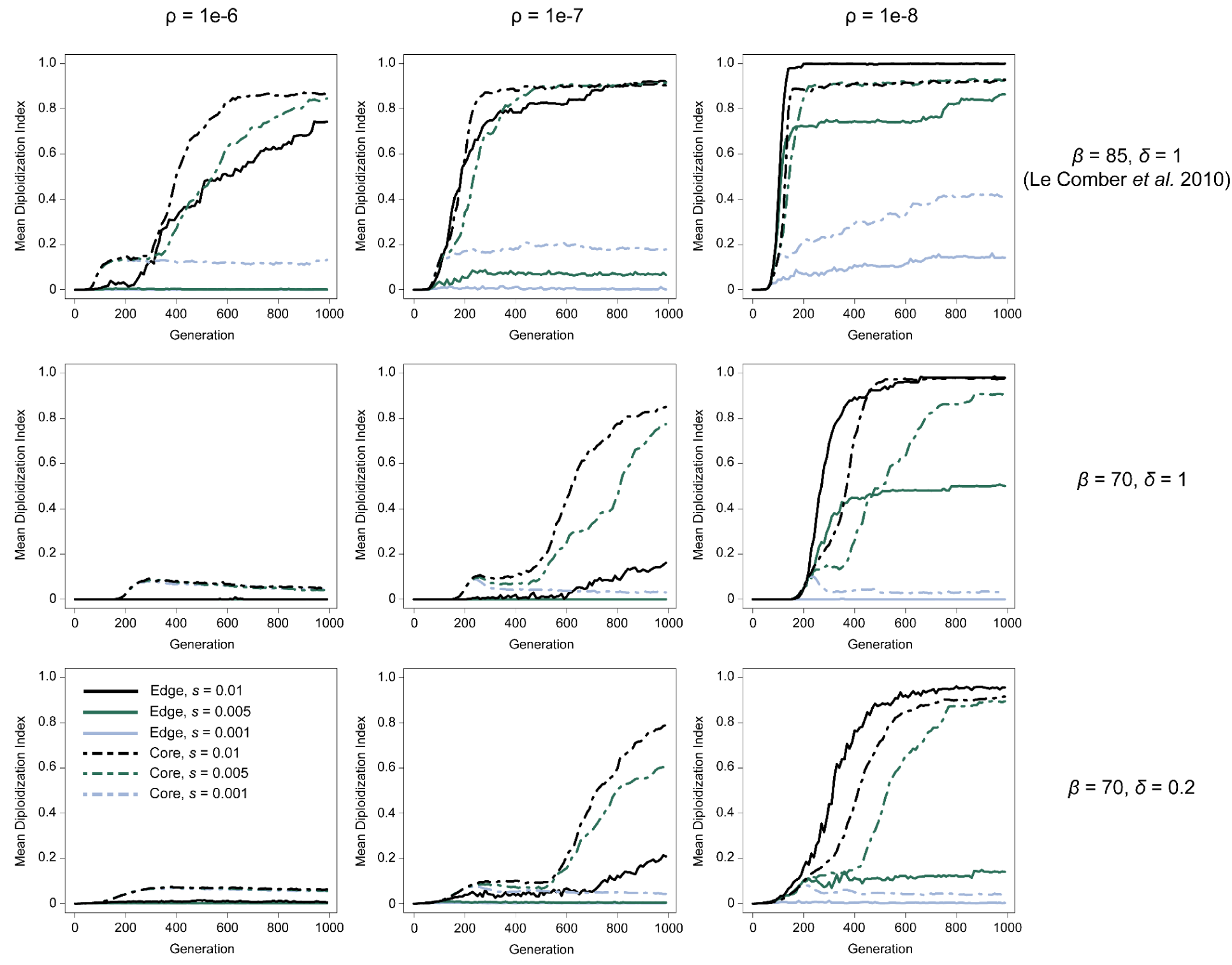
Diploidization of core and edge populations under the pairing efficiency model at *λ* = 1000. Lines represent the mean diploidization index (out of 50 for each model) in the edge (solid lines) or core (dashed lines). Line color delineates the selective coefficient (*s*), and individual figures are separated by recombination rate (⍴; columns) and the parameter set for the model full disomy (*β* and δ; rows).

**Supplemental Figure 13.**
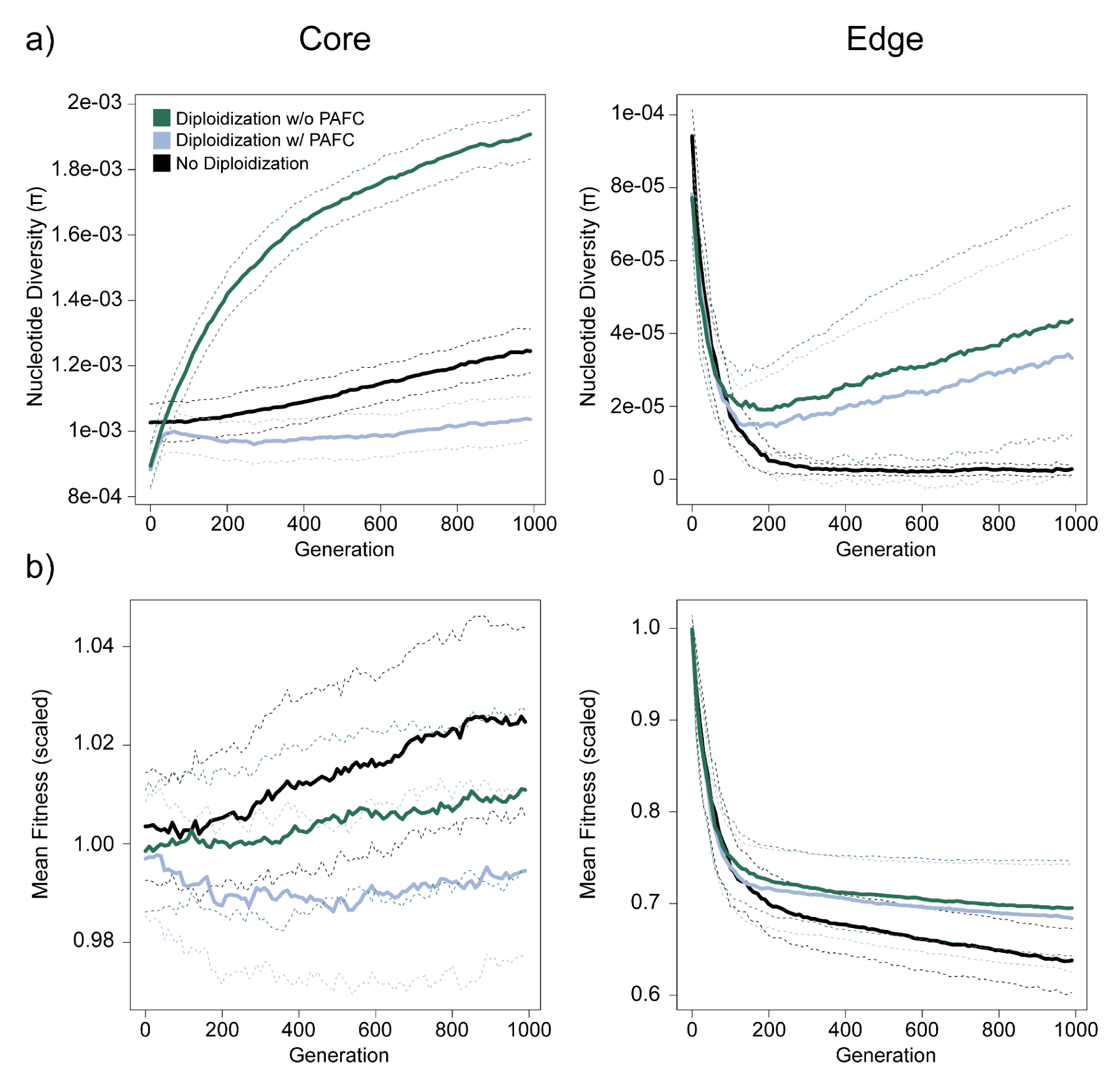
The cost of pairing efficiency regulated diploidization on fitness and nucleotide diversity. a) Nucleotide diversity (π) in each generation of core and edge populations when the model includes no diploidization, diploidization under the pairing efficiency model with pairing associated fitness costs (PAFC), and diploidization under the pairing efficiency model without PAFC. b) Mean fitness (scaled at the start of expansion) in each generation of core and edge populations when the model includes no diploidization, diploidization under the pairing efficiency model with pairing associated fitness costs (PAFC), and diploidization under the pairing efficiency model without PAFC. All simulations run with *λ* = 100, *β* = 85 δ = 1, *s* = -0.005, and ⍴ = 1×10^-6^. Note the different *y*-axis scales for the edge and core populations.

**Supplemental Figure 14.**
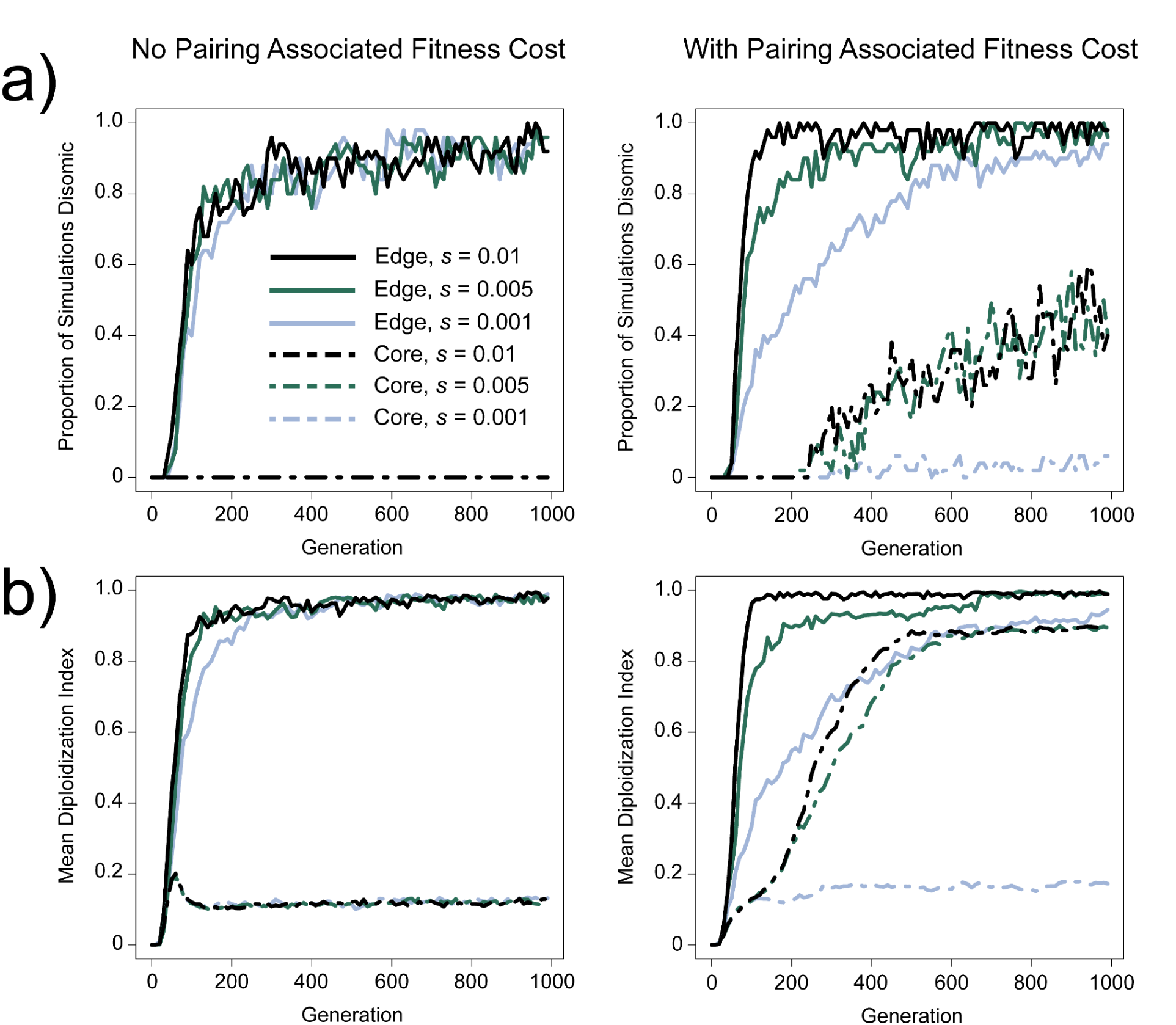
Interaction of selection coefficient of standard deleterious mutations and the fitness of gametes determined by pairing efficiency on the evolution of disomy under the pairing efficiency model of diploidization. a) The proportion of simulations where disomy (diploidization index ≥ 0.9) has evolved if pairing efficiency affects (right) or does not (left) affect gamete fitness. b) Average diploidization index for each simulation if pairing efficiency affects (right) or does not affect (left) gamete fitness. For each parameter 50 simulations were run under a model where *λ* = 100, *β* = 85, δ = 1, *s* = 0.005, and ⍴ = 1e-8.

**Supplemental Figure 15.**
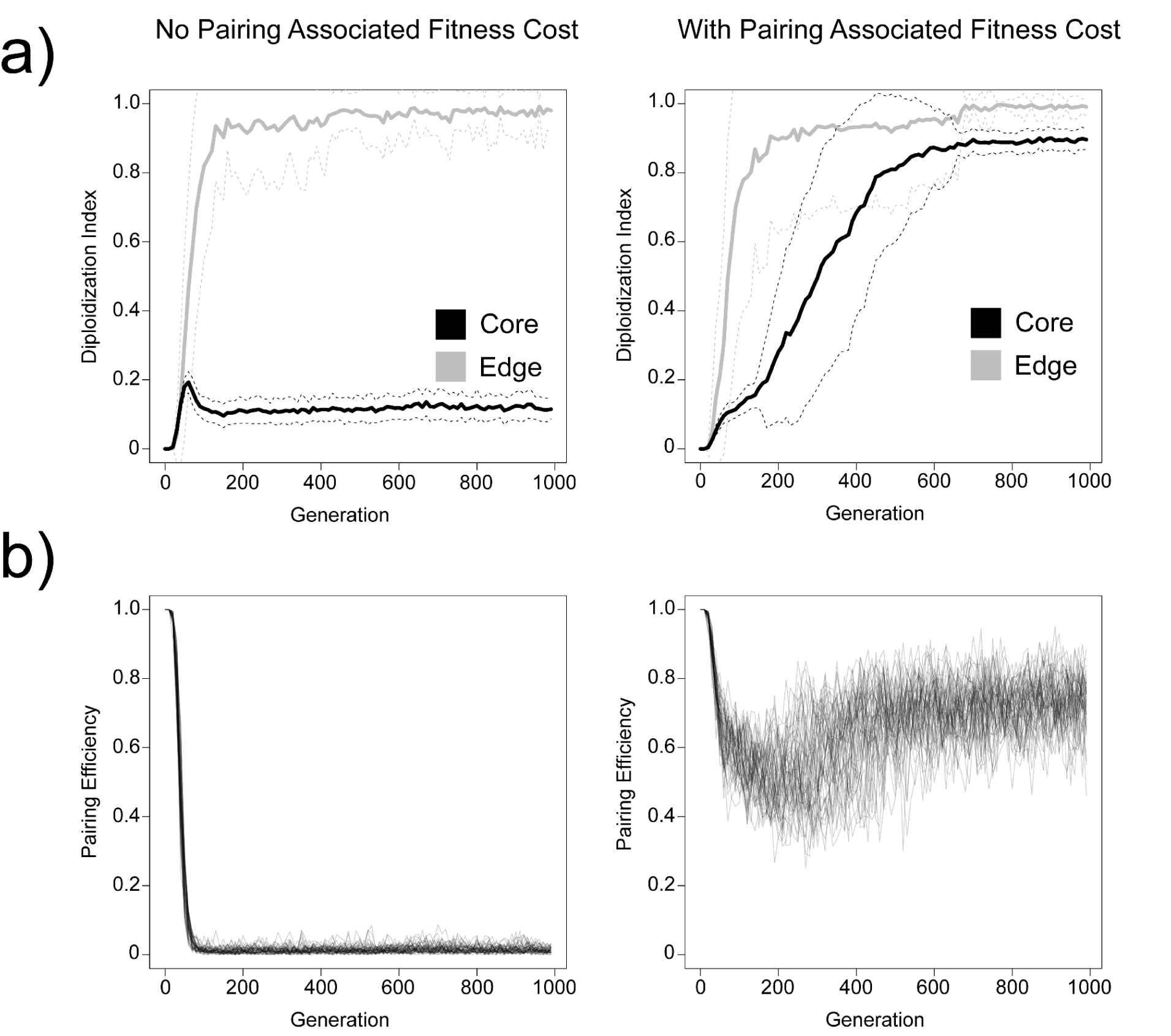
Effect of pairing efficiency affecting gamete fitness in core populations where disomy does evolve. For all figures, *λ* = 100, *β* = 85, δ = 1, *s* = 0.005, and ⍴ = 1×10^-6^. a) Average diploidization index for each simulation if pairing efficiency affects (right) or does not affect (left) gamete fitness. Dashed line represents ± 1 standard deviation, averages and standard deviation calculated from 50 independent simulations. b) Pairing efficiency of core populations if pairing efficiency affects (right) or does not affect (left) gamete fitness. Each line represents a single simulation.

**Supplemental Figure 16.**
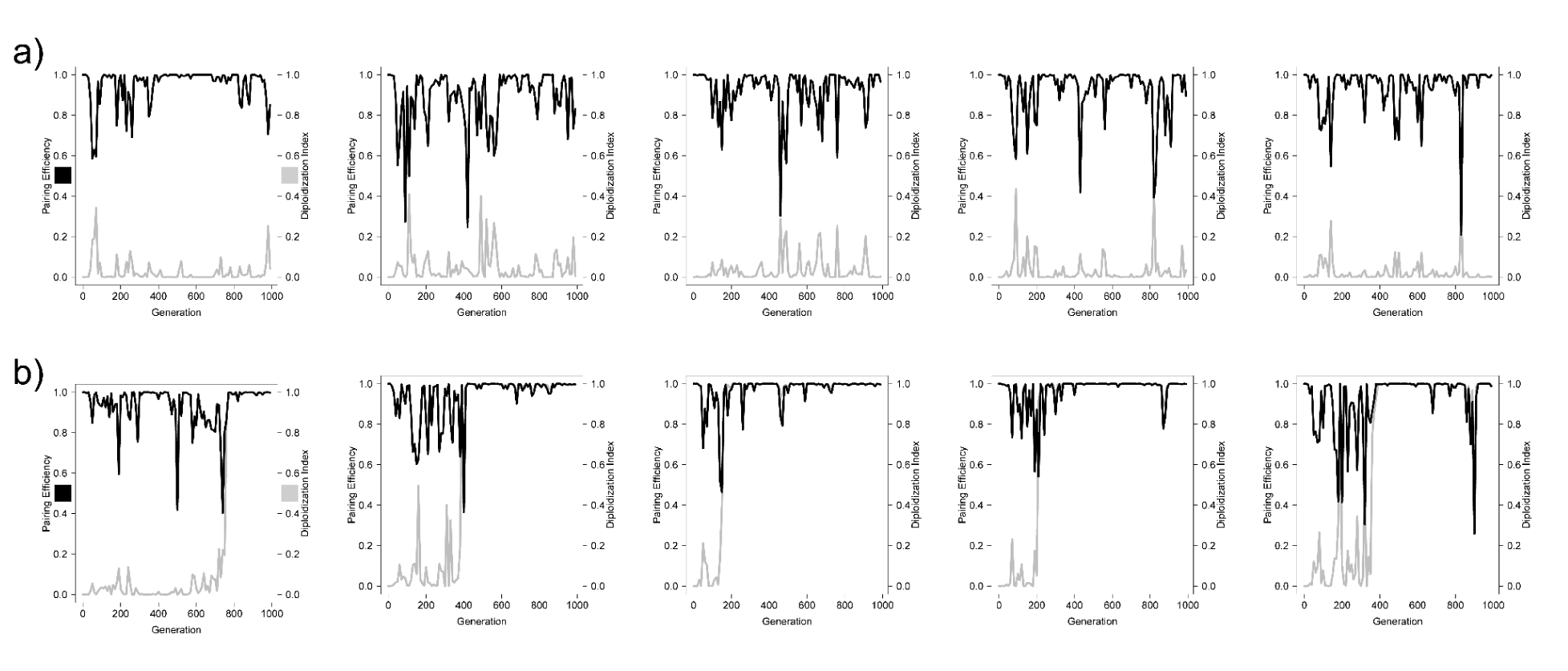
Pairing efficiency (black) and diploidization index (gray) of edge populations when disomy did not (a) or did (b) evolve within 1000 generations. Each figure represents a single simulation under a model where *λ* = 100, *β* = 85, δ = 1, *s* = 0.005, and ⍴ = 1e-6.

**Supplemental Figure 17.**
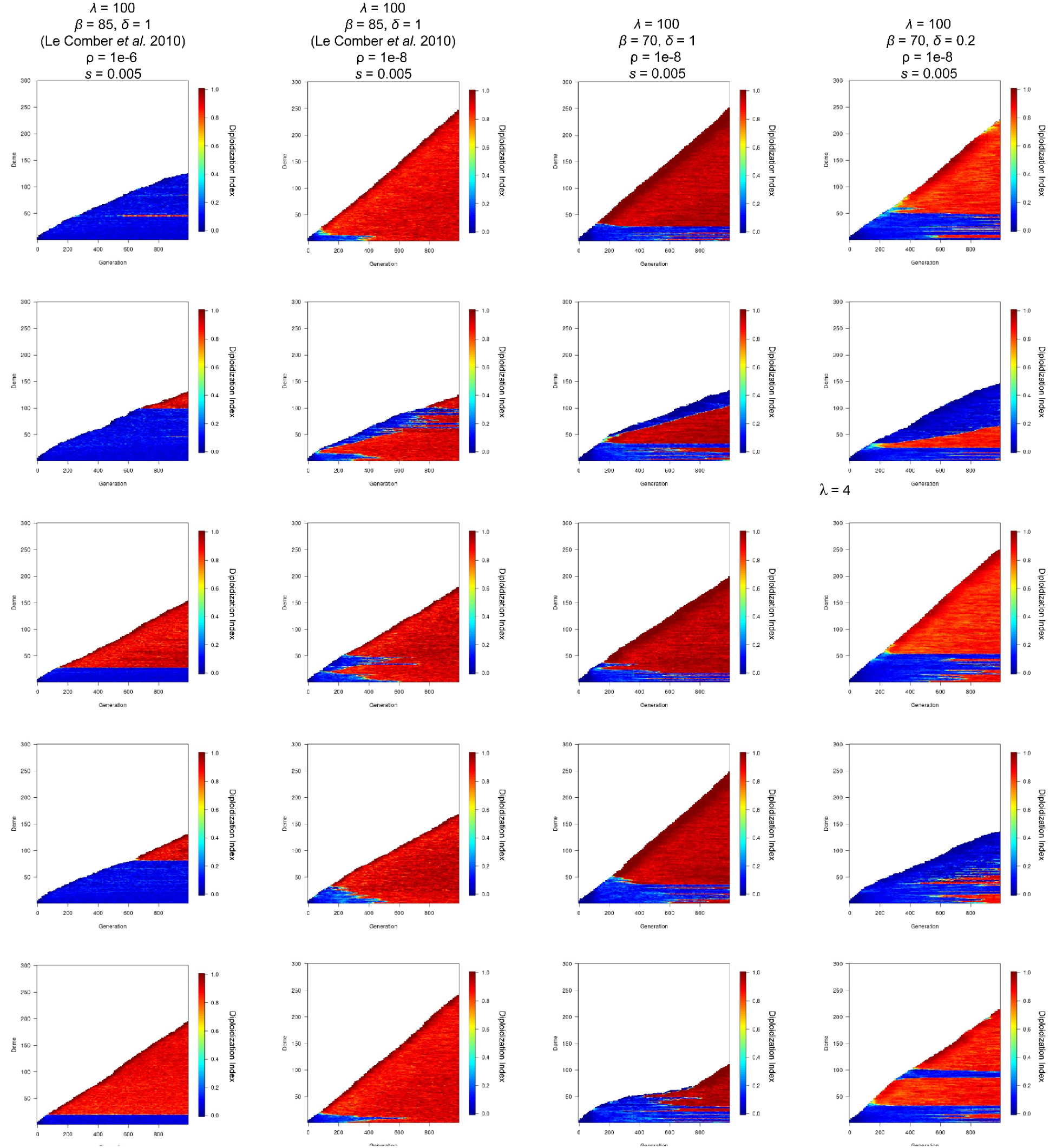
Diploidization index of example individual simulations under the pairing efficiency diploidization model for *λ* = 100.

**Supplemental Figure 18.**
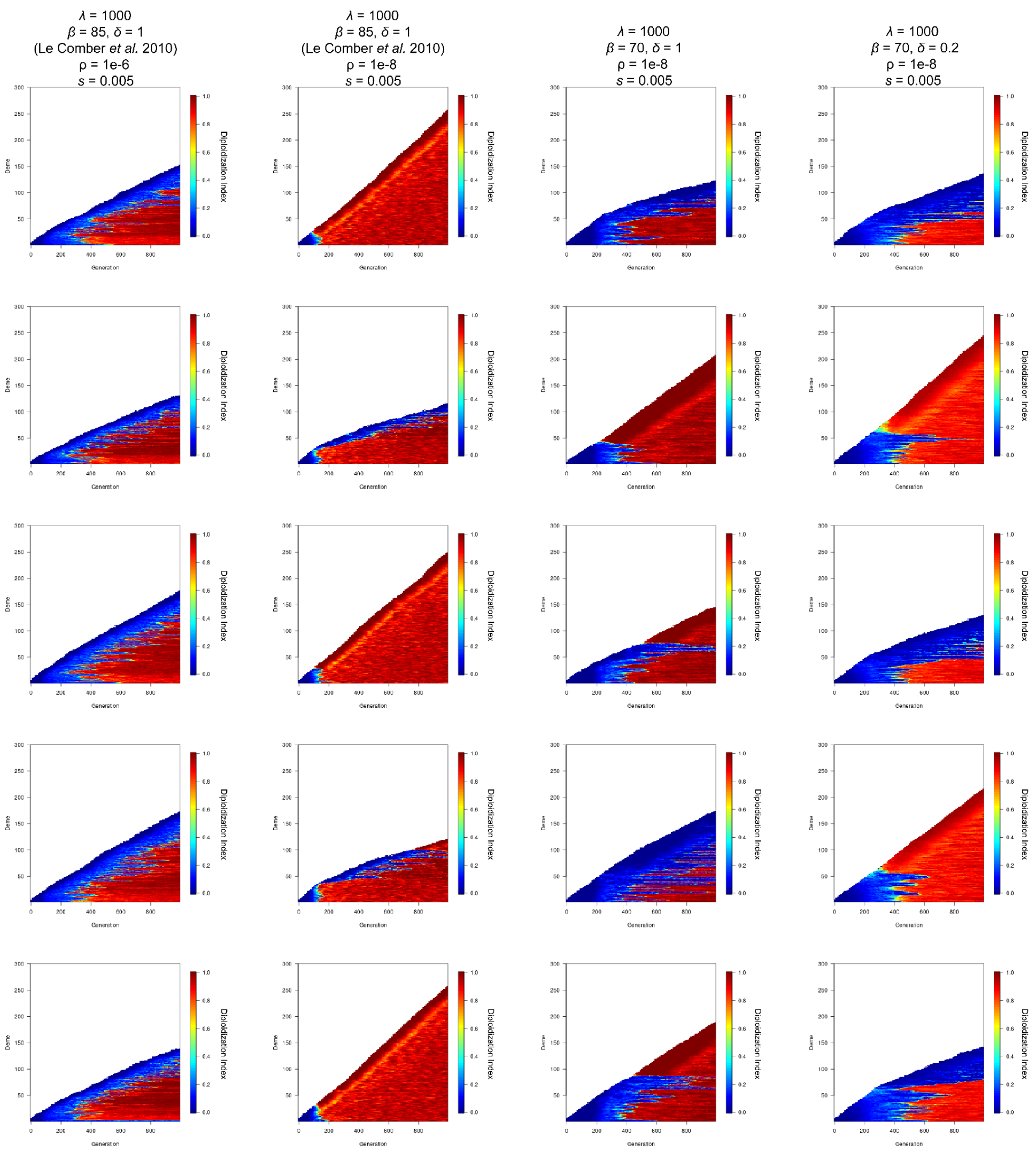
Diploidization index of example individual simulations under the pairing efficiency diploidization model for *λ* = 1000.

**Supplemental Figure 19.**
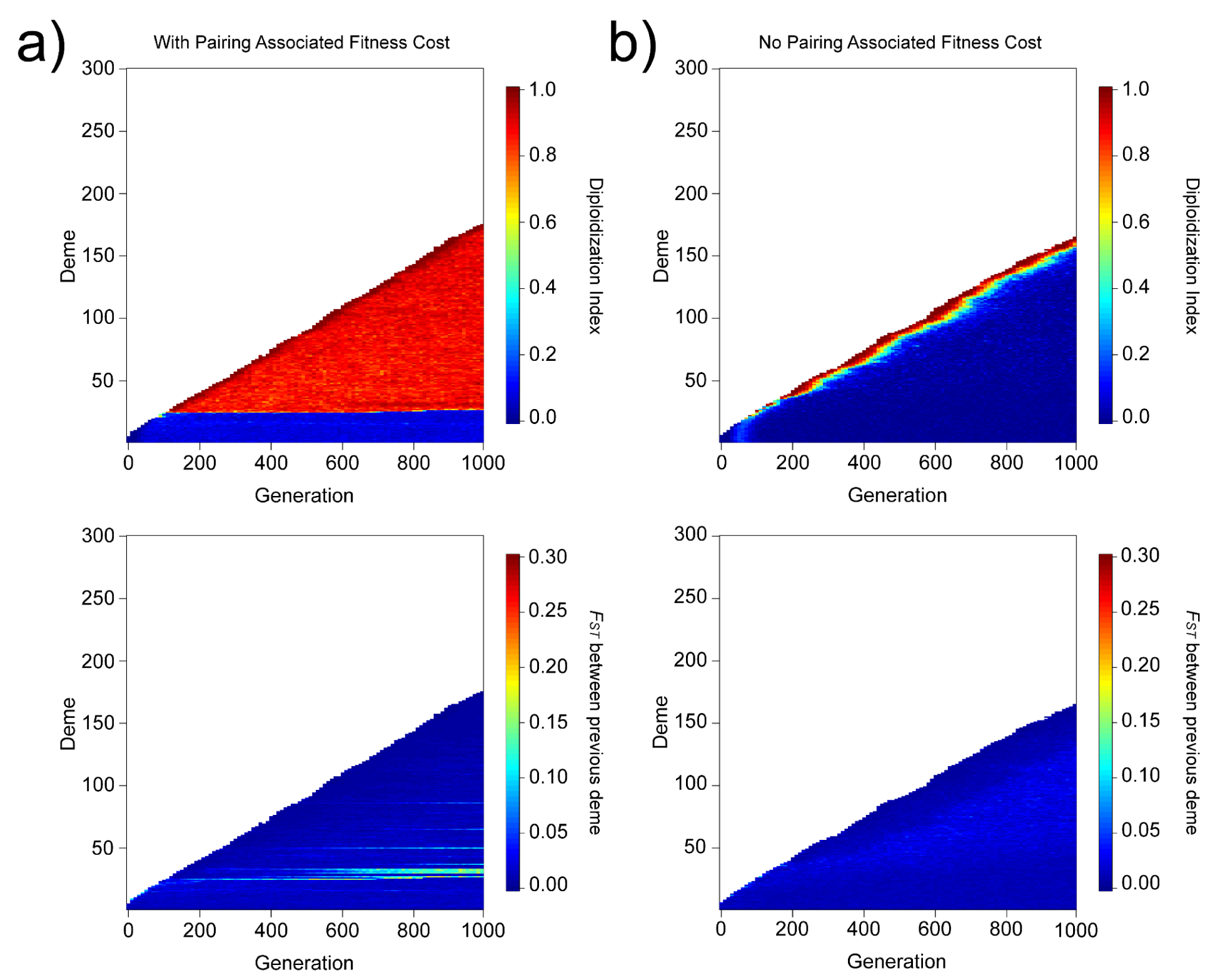
Pairing associated fitness costs maintain disomy once evolved. a) Diploidization index (top) and F_ST_ between focal and previous deme (bottom) when diploidization involves pairing associated fitness costs. b) Diploidization index (top) and F_ST_ between focal and previous deme (bottom) when pairing associated fitness costs are not included in the diploidization model. All simulations run with *λ* = 100, *β* = 85 δ = 1, *s* = 0.005, and ⍴ = 1×10^-6^.

